# FreiPose: A Deep Learning Framework for Precise Animal Motion Capture in 3D Spaces

**DOI:** 10.1101/2020.02.27.967620

**Authors:** Christian Zimmermann, Artur Schneider, Mansour Alyahyay, Thomas Brox, Ilka Diester

## Abstract

The increasing awareness of the impact of spontaneous movements on neuronal activity has raised the need to track behavior. We present FreiPose, a versatile learning-based framework to directly capture 3D motion of freely definable points with high precision (median error < 3.5% body length, 41.9% improvement compared to state-of-the-art) and high reliability (82.8% keypoints within < 5% body length error boundary, 72.0% improvement). The versatility of FreiPose is demonstrated in two experiments: (1) By tracking freely moving rats with simultaneous electrophysiological recordings in motor cortex, we identified neuronal tuning to behavioral states and individual paw trajectories. (2) We inferred time points of optogenetic stimulation in rat motor cortex from the measured pose across individuals and attributed the stimulation effect automatically to body parts. The versatility and accuracy of FreiPose open up new possibilities for quantifying behavior of freely moving animals and may lead to new ways of setting up experiments.

## 1 Introduction

Systems neuroscience strives to assign functions to neuronal circuits and their activity. Often, these functions are defined by behavior in a task carefully designed to cover specific aspects, e.g. rule learning, working memory or reaching to a defined target. However, neuronal activity can be dominated by uninstructed movements not required for the task. Confusingly, some uninstructed movements can be aligned to trial events while other movements might occur idiosyncratically, accounting for trial-by-trial fluctuations that are often considered ‘noise’ [1]. This is even true for head-fixed animals [2, 3]. In order to address this problem, detailed tracking of the animals’ movements including single body parts and correlating them to neuronal activity is essential. Several new tools are already available for interpreting video data. However, existing tools are limited by one of several aspects: (1) Marker-based approaches [4] influence natural movements, are restricted to applicable body sites and rely on the tolerance of the animal. (2) So far, marker-free analyses have only been applied in 2D, [5–7], thus preventing the true pose reconstruction of freely moving animals covering all three dimensions of their movements. By using multiple cameras for video recording, 2D outputs can be triangulated to yield 3D reconstructions a posteriori [8, 9]. However, such post-processing suffers from the ambiguities in the initial 2D analysis reducing accuracy and reliability. Indeed, exactly this tracking accuracy is crucial to subdivide movements into well-defined trajectories and behavioral categories for isolated analysis.

Here, we introduce the new tracking tool FreiPose, which allows reconstructing detailed body poses and single body part movements directly in 3D. Based on synchronized multi-view videos, FreiPose calculates a sparse reconstruction of on-body keypoints. These keypoints are freely definable in terms of number, position, and semantic meaning (e.g. eye, paw etc.). FreiPose operates in a tracking-by-detection fashion, in which each time step is processed independently. We demonstrate that FreiPose is a versatile tool which is easily adaptable to different applications, e.g. tracking of various organisms, full-body tracking or detailed tracking of paws including individual digits. By combining this tool with extracellular laminar recordings in the rat motor cortex, we recapitulated that a large fraction of neurons (42.5%) was tuned to body poses, as has previously been shown with a marker-based approach [4]. Our detailed tracking enabled automatic clustering of movements into intuitively meaningful behavioral categories. Applying this behavioral categorization in freely moving rats, we found clear neuronal tuning to single paw trajectories similar to what has been described previously in head fixed animals involved in constrained tasks [10, 11]. Moreover, we used FreiPose to quantify the effect of optogenetic stimulation of the motor cortex based on the rats’ movements. Importantly, FreiPose allowed attributing the stimulation effect to individual body parts as well as a temporal analysis of the stimulation effect. In summary, FreiPose is particularly suited for studies conducting physiological recordings in freely moving animals. It allows conclusions about the behavioral state in trained as well as in spontaneously behaving animals including detailed information of individual body parts.

## 2 Results

Given an experimental setup with multiple cameras (Fig. 1**a**), FreiPose leverages a novel 3D convolutional network architecture to reconstruct 3D poses by integrating information across all available views (Fig. 1**b**) instead of treating cameras separately (Fig. 1**c**). It calculates the 3D positions of predefined key-points on the animal’s body. FreiPose is easily adaptable to other relevant applications, e. g. pose reconstruction of mice (Fig. 1**f**) or pellet reaching (Fig. 1**g**). The system can be tailored to an arbitrary type of animal or different set of 3D keypoints, by following an iterative three-step reconstruct-inspect-refine scheme (Fig. Sup. 1 for workflow overview). Using a synchronized and calibrated video sequence, the pre-trained network model yields pose reconstructions. Guided by reconstruction confidence, potentially erroneous frames can be identified quickly and are selected for manual correction in a human-in-the-loop approach (see Fig. Sup. 3**a**). For a completely new experimental setup or set of keypoints, the guidance by reconstruction confidence is initially omitted and for bootstrapping a small number of frames is sampled uniformly from the videos instead. Multiple persons can annotate videos in parallel using a stand-alone, intuitive drag-and-drop GUI that leverages multi-view constraints during annotation (see Fig. Sup. 3**b**). The method is retrained with the additional annotation, leading to pose reconstructions with fewer and smaller errors. FreiPose comes with a *Docker* installation package, which simplifies the software installation, and tutorials with exemplary data and step-by-step videos. All necessary information for applying FreiPose can be found in the released Github repositories.

**Figure 1:**
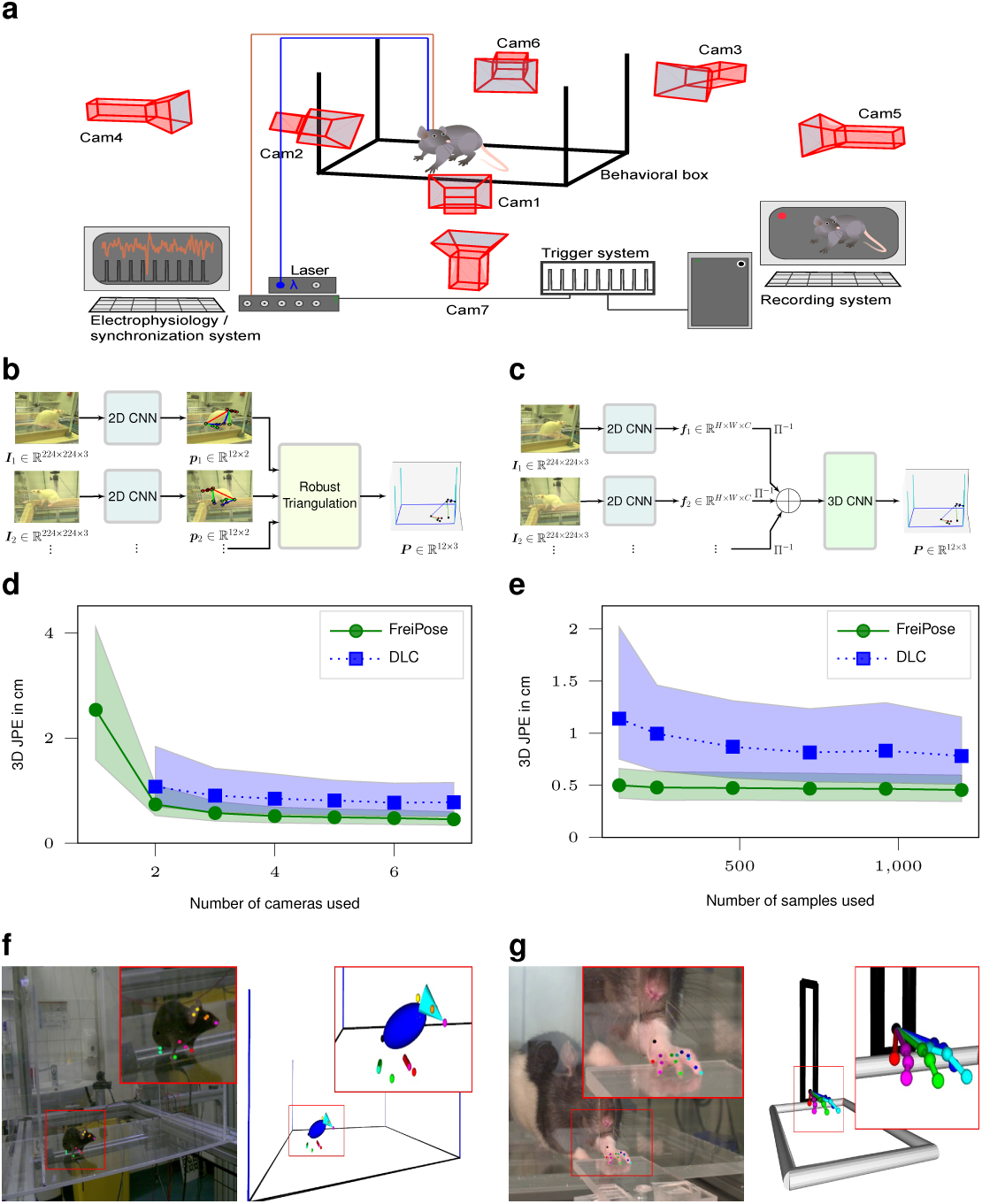
Overview over the proposed motion capture framework and its evaluation. **a** Motion capture setup with required hardware elements. Orange line - connection for electrophysiology, blue line - fiber optics for optogenetic stimulation. **b** State-of-the-art motion capture methods reconstruct independent 2D poses for each view independently and subsequently calculate a 3D pose, which requires resolving ambiguities in each view separately. **c** FreiPose accumulates evidence from all views leading to a holistic 3D pose reconstruction. Ambiguities are resolved after information from all views is available. **d, e** Median of the Joint Position Error in cm for different number of cameras and number of samples compared to the DeepLabCut (DLC) [6] based single view network. Results are based on 3 training trials on an evaluation set with 614 samples. Shaded areas refer to the 30% and 70% percentiles. FreiPose is more data efficient and performs better regardless of the number of cameras. Largest differences can be observed for highly articulated keypoints, e. g. the front paws (see Fig. Sup. 4). **f, g** FreiPose can be easily adapted towards other tasks, e. g. pose reconstruction of mice or paw reconstruction including individual digits during a pellet reaching task (see Fig. Sup. 5 and supplemental videos 1-3).

### High motion capture accuracy and reliability

We measured the accuracy (in terms of median 3D error) and reliability (in terms of percentage of samples below a maximum error bound) of FreiPose on video recordings of freely moving rats consisting of 1813 manually labeled samples with 12 distinct body keypoints. The frames are sampled from 12 recording sessions featuring 5 different individuals (3 Long-Evans (hooded) and 2 Sprague Dawley (albino) rats). Some animals were recorded once, others on several days. We split the recordings into training and evaluation set, resulting in 1199 training samples and 614 evaluation samples. Each sample contains 7 images recorded simultaneously from different cameras simultaneously and a single manually annotated 3D pose consisting of a 3D location for each keypoint which is obtained from manual 2D annotation in at least two camera views.

For comparison, we trained DeepLabCut (DLC) [6], a popular tool for 2D keypoint tracking, on the same dataset of images and applied standard triangulation methods to yield 3D poses [9]. FreiPose compares favorably in terms of the number of camera views required to reach a certain accuracy (Fig. 1**d**), data efficiency (lower median error with the same number of labeled samples, Fig. 1**e**), accuracy (median error of 4.54 mm vs. 7.81 mm for the full sample setting, Fig. 1**e**), and reliability (percentage of keypoints with an error smaller than 7.5 mm is 82.8% vs. 48.1%, Fig. Sup. 4**b**).

### Body pose reconstruction recapitulates neuronal tuning

To evaluate the applicability of FreiPose for experimental measurements, we extracted single paw movements as well as body and head pitches (Fig. 2**a**) from FreiPoses’ 3D reconstructions and combined them with extracellular recordings in the rat motor cortex. Previous work on the neuronal representation of body poses in rodents relied on marker-based systems [4, 12]. Applying FreiPose to one recording session of a freely exploring rat combined with simultaneous neural recordings revealed that almost half of the recorded neurons in motor cortex was tuned to postural features (42.4%, 28/66 cells, Fig. 2**b**). The neuronal pose representation was confirmed in 2 additional rats. Neurons were considered tuned, if the z-score exceeded a critical value (corresponding to a Bonferroni corrected p-value < 0.05) relative to the shuffled distribution for at least 3 consecutive postural bins. This fraction of tuned neurons is in accordance with the previously reported percent of cells in motor cortex based on the use of reflective markers [4]. This demonstrates that FreiPose is able to effectively capture the poses with a precision which allows claims about neuronal correlations.

**Figure 2:**
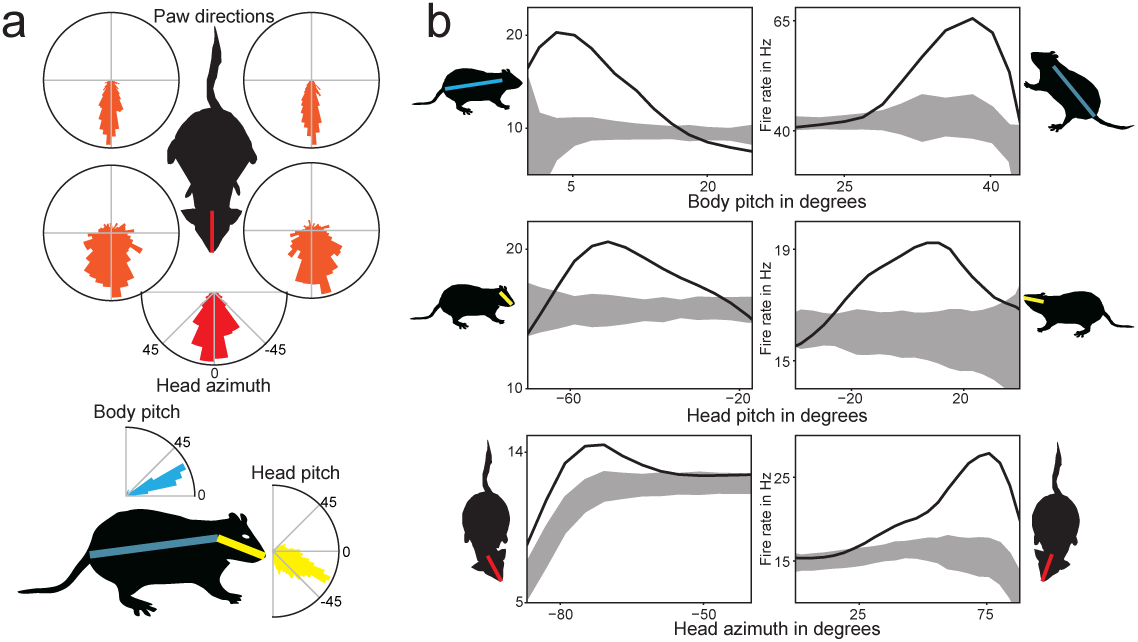
FreiPose recapitulates known neuronal pose representations in rat motor cortex. **a** Circular plots of movement distributions over directions for each paw and the head. **b** 1D tuning curves of representative neurons for body pose variables (e.g. body and head pitch as well as head azimuth) contrasted by randomly time-shifted data. Gray shading indicates standard deviation of shuffled distributions. Rat pictographs refer to the preferred pose of the respective neuron.

### Body poses reveal behavioral categories

The output of FreiPose is used to split the videos into segments according to behavioral categories. To this end, we modify an unsupervised behavioral clustering algorithm [13] to seamlessly work with pose reconstructions results from FreiPose. Based on the reconstruction results, distances of all keypoints relative to each other are calculated as well as to the floor. From these distances, main components via PCA are extracted on which wavelet based spectrograms representing postural changes are computed. We combined the information about the current pose (body-centered keypoint locations) with the postural changes (PCA based spectrograms) into one matrix and embedded both sets of features non-linearly into 2D space using t-SNE [14]. This embedding preserves local relationships. Thus, neighboring points in the embedding relate to similar postural components, putatively referring to congruent behaviors. Individual high density clusters are separated by a watershed algorithm. Afterwards, video segments based on the clustering are used to manually assign a behavioral category to each cluster (Fig. Sup. 6, *Supp_Video_4_BehaviorClusters*.*avi*).

We obtained a density map of behavior (Fig. 3**a**) and identified 4 clearly distinguishable behavioral clusters: locomotion, cleaning, rearing, and inspection of nearby environment, i.e. small sideway movements. By defining intuitively meaningful variables (i.e. body pitch, average paw speed, paw distance to nose and horizontal body speed) we are able to confirm these behavioral categories (Fig. 3**c**); e.g., during cleaning, the distance between paws and nose was lowest. This method can identify the behavioral state of an animal either during spontaneous movements or during instructed tasks [1].

**Figure 3:**
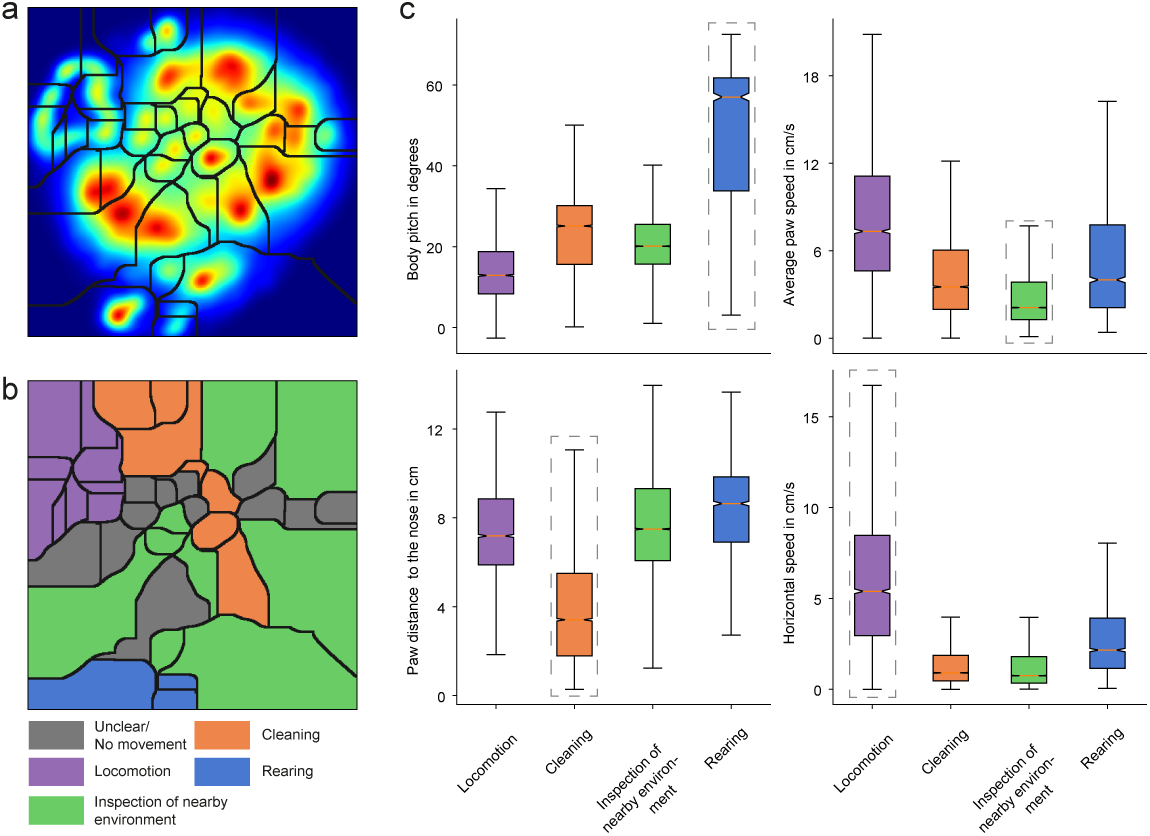
Unsupervised clustering of behavior. **a** Density estimate of the non-linear embedding and watershed clustering. **b** Cluster identities based on visual inspection of the behavior. **c** Quantitative analysis of behavior in each cluster. For each of the combined clusters, a variable is identified which intuitively describes the cluster (gray dashed rectangle). All distributions are significantly different (two sample Kolmogorov–Smirnov test, Bonferroni adjusted p < 7*e -* 13).

### Neuronal representations of behavioral states and paw trajectories

To test whether we can identify a systematic neuronal representation of spontaneous movements outside of the classical instructed tasks, we extracted single paw trajectories. To this end, we divided continuous paw movements during exploratory behavior into artificial trials by setting a threshold of the paw speed (for details see methods 6.3.4). In line with previous work based on instructed tasks [15], neurons in motor cortex were modulated by the movements of the contralateral but not the ipsilateral paw in a ± 333 ms window around movement onset (Fig. 4**a,b**, two-sided t-test on mean normalized firing rate across population, p< 0.05). At the individual neuron level, 12% (8/66 neurons) showed a significant difference (one-way ANOVA per neuron Bonferroni adjusted p< 0.05). In contrast, the reconstructions via DLC did not allow the detection of a significant neuronal bias towards one paw (Fig. 4**b**) underlining the importance of holistic 3D pose reconstruction.

**Figure 4:**
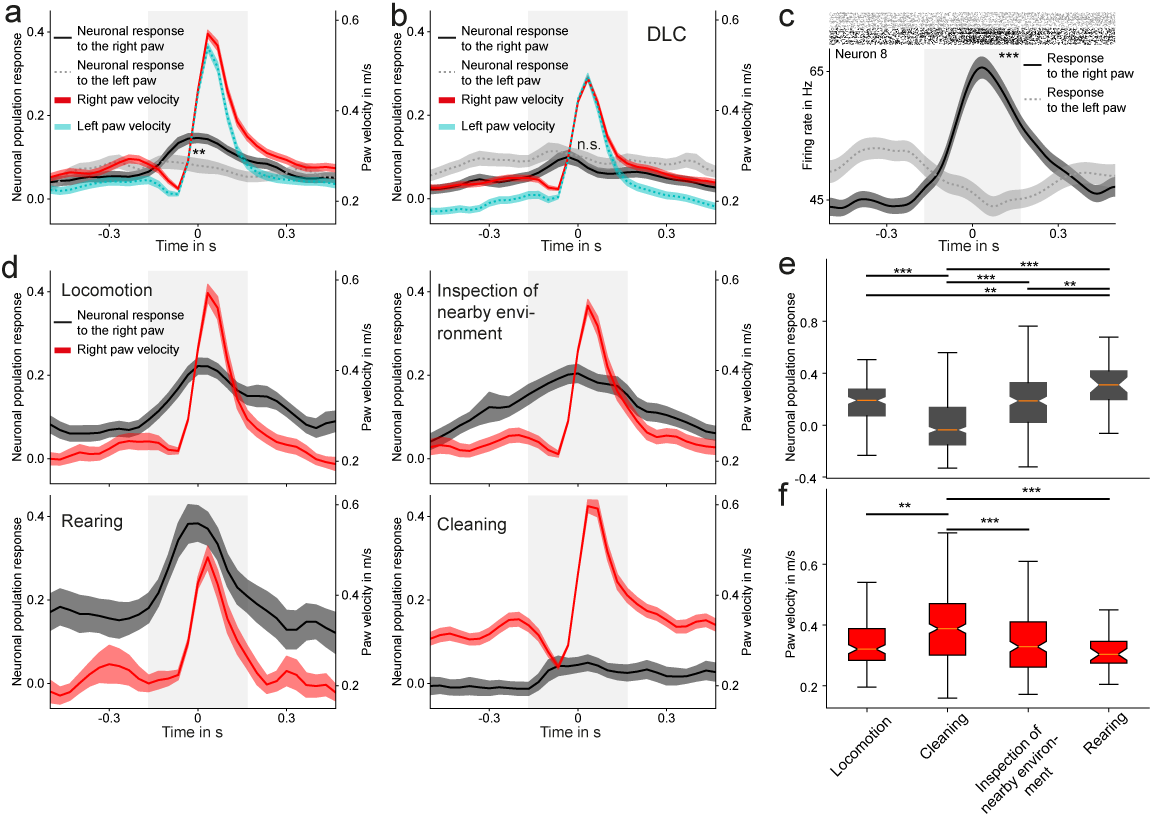
Behavioral clustering reveals differential neuronal tuning to contralateral paw movements. **a** Motor cortex neurons responded to the movements of the right paw (contralateral to the electrode implantation) differently than to the left paw. **b** DLC was not able to recapitulate the significantly different tuning to left and right paw movements. **c** FreiPose was able to identify side specific tuning on a single neuron level. The example neuron was significantly tuned to movements of the right paw. **d** Population responses (gray) to the movement of the right paw during different behaviors with corresponding paw velocities (red). **e** Summary of the population responses to the movement of the right paw during different behavioral categories. **f** Summary of the velocities of the right paw during different behavioral categories. The notch represents the 95% confidence interval around the median, whiskers refer to 1.5 times of the interquartile range. Outliers were omitted for visualization purposes. Solid lines - mean, shaded areas - standard error. ** indicates p< 0.01, *** p< 0.001 two-sided t-test, Bonferroni adjusted.

To test whether different behavioral categories had an influence on neuronal representations of paw movements [16], we isolated behavioral categories and reanalyzed the neuronal responses. Indeed, neurons responded significantly different between the behavioral categories (except between locomotion and exploration) to the movements of the contralateral paw. However, the velocity of the paw movements differed only significantly during cleaning (one-way ANOVA p< 0.05, post-hoc Bonferroni adjusted two-sided t-test, Fig. 4**d,e,f**), thus ruling out a simple velocity tuning of the neurons.

Using the precise 3D tracking of FreiPose we analyzed individual paw trajectories of the two behavioral categories “locomotion” and “inspection of nearby environment” as these two categories contained step-like movements. We asked whether neurons responded differently to similar movement trajectories if those occurred in a different behavioral context. Indeed, when we compared the response of individual neurons to paw movements during those behavioral categories, we found neurons with significantly different responses (10.6%, 7*/*66, one-way ANOVA per neuron Bonferroni adjusted p< 0.05, Fig. 5 **a,b**). To investigate whether the neuronal differences relied on the exact paw trajectories or on the behavioral state, we clustered paw trajectories by k-means. When adjusting for the occurred trajectories during both behaviors (i.e. taking only similar trajectories into account), no significant difference was observed (Fig. 5 **c,d**). Additionally, we found individual neurons with significant modulation of their response relative to the trajectory class (16.7%, 11/66, one-way ANOVA, Bonferroni adjusted p< 0.05, Fig. 5 **e,f**). This suggest that neurons in motor cortex code for the movement trajectory of the paw, independent of the here investigated behavioral contexts in which those movements occurred. Applying the pose reconstructions of DLC impaired the detection of neuronal response differences (Fig. Sup. 7).

**Figure 5:**
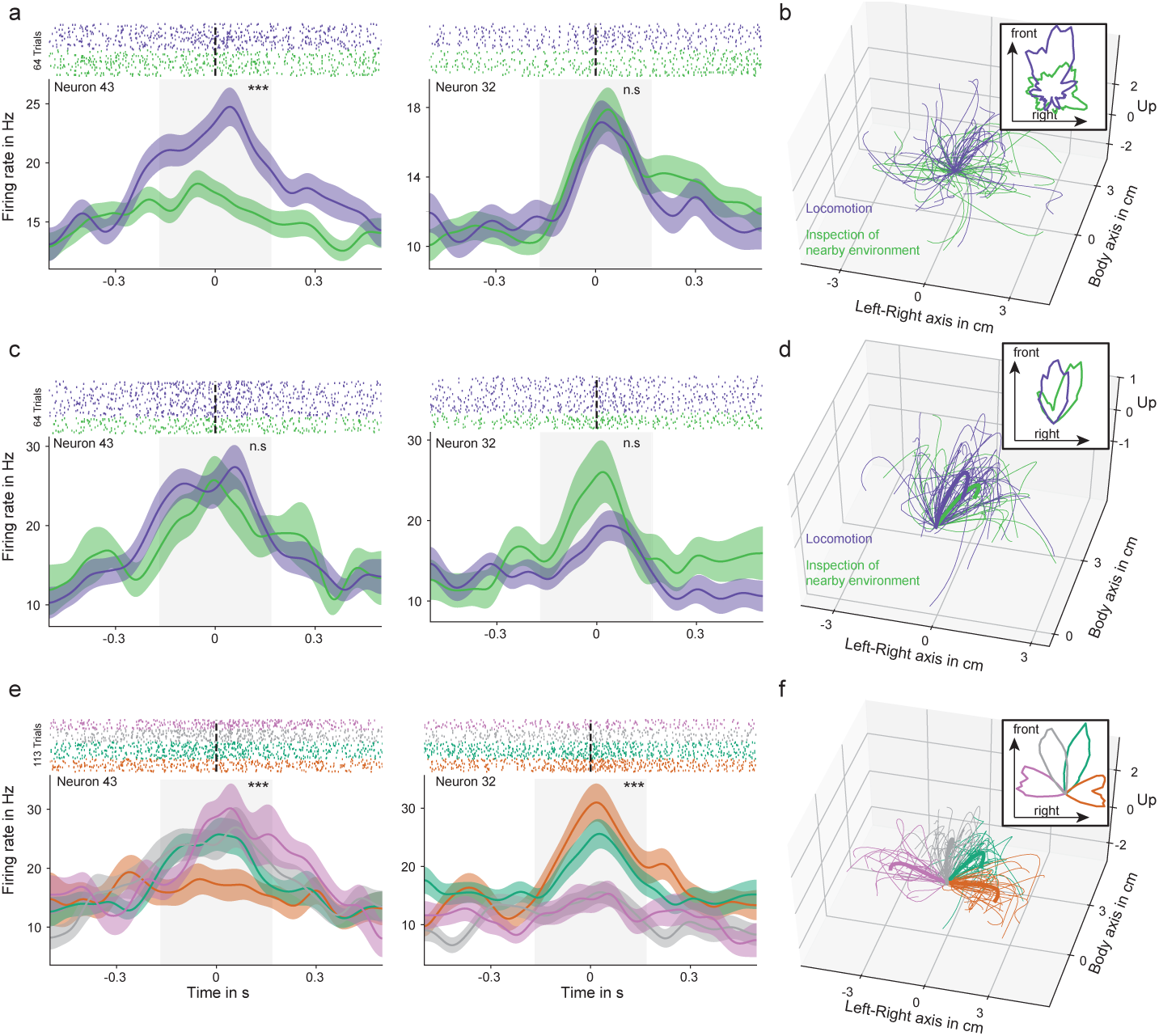
Direction selective neuronal representation of paw trajectories is independent of behavioral context. **a** Two example neurons with a significant response to movements of the right paw. Neuron 43 (left panel) discriminated significantly between behavioral states (blue - locomotion, green - inspection of nearby environment) while neuron 32 (right panel) did not discriminate. For visualization purposes, the raster plots contain a random subset of trials. **b** Trajectories of the right (contralateral) paw during locomotion and inspection of nearby environment. A subset of trajectories is shown for visualization purposes. **c** The comparison of neuronal responses to subsets of directional similar trajectories during different behavioral clusters reveals no significant differences. **d** Directionally similar trajectories of the right paw classified according to behavioral state (subset from panel b, corresponding to the combination of gray and green trajectories in panel f). **e** Example neurons with significant modulation and tuning preferences for subsets of trajectories. **f** Paw trajectories during step-like paw movements (based on the behavioral clusters “locomotion” and “inspection of nearby environment”). We focused on anterior movements (four clusters of trajectories corresponding to the 9 to 3 o’clock directions). Color represent the investigated four paw directions in the rats’ body reference frame. Inserts represent the distributions of the horizontal projections of movement directions. *** indicates p < 0.001, one-way ANOVA, Bonferroni adjusted (see Fig. Sup. 7 for a comparison to DLC).

### Quantifying the effect of optogenetic stimulation

Manipulations in motor cortex are likely to impact movements. This effect can be quantified via coarse measurements, e.g. rotational behavior, speed, mobile time, mobile episodes, and distance traveled [17, 18]. Recently, effects on single body parts have also been started to be investigated via video analysis [19]. To systematically investigate the effect of optogenetic stimulation in freely moving rats, we recorded the movements of three animals in four sessions, two sessions with 30 Hz laser burst frequency and two sessions with 10 Hz with a stimulation duration of 5 sec, 10% duty cycle. During each recording session we stimulated each animal 5 to 7 times, with a minimum inter-stimulus time interval of 45 sec. We retrained FreiPose based on 136 samples from these recordings and systematically defined 908 behavioral variables for every time step from the reconstructed poses. Behavioral variables included transformation of the pose into a rat-aligned Cartesian coordinate frame, the distance of keypoints with respect to the ground floor as well as their velocities and Fourier transforms. Additionally, we calculated angles between body limbs with respect to each other and the direction of gravity (e. g., angle between head and body axis, see Table Sup. 2 for the full list of variables).

To reveal changes in behavior, we followed an *attribution-by-classification* paradigm: given the behavioral variables at a time step *t* we trained a linear SVM model (*C* = 0.0025) to classify every time step into stimulated (i. e., *positive*) or not stimulated (i. e., *negative*). We trained separate classifiers for each animal, used one recording for training and left one for evaluation. The resulting classifiers achieved a balanced average accuracy of 59.1% to 73.1% on their evaluation sets. The average classifier response was able to follow the temporal dynamics of the stimulation effect and revealed an increasing effect over the course of stimulation (Fig. 6**a**).

**Figure 6:**
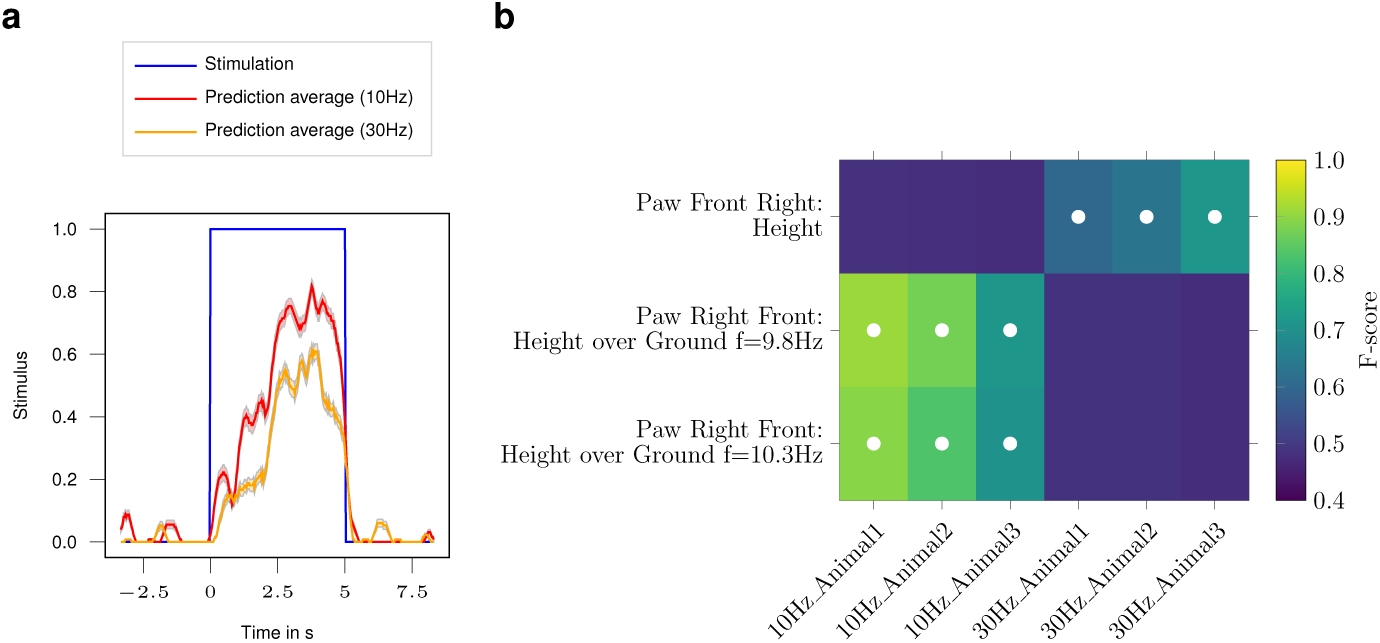
Automatic evaluation of the optogenetic stimulation effect via FreiPose. **a** Automatic detection of the behavioral effect during optogenetic stimulation with temporal resolution. The classifier predicted whether a time step was stimulated ‘1’ or not ‘0’ based on the calculated behavioral variables. We averaged the predicted stimulus of the classifier model within temporally aligned windows across stimulation trials on recordings which were withheld for evaluation (18 and 19 trials for the 10 Hz and 30 Hz, respectively). The stimulation spans from 0 sec to 5 sec and the prediction scores trend indicates an increasingly visible effect over time. Curves are temporal smoothed averaging over a window length of 23 ms. Shaded areas indicate the Standard Error of the Mean. **b** Attribution of the stimulation effect to individual body parts. We trained an ensemble of classifiers to distinguish stimulated from non stimulated frames given only a single behavior variable as input to each classifier. Analyzing the resulting classifiers allowed to distinguish important factors from less important ones. The color scale refers to the *F-score* of the respective classifier and white dots indicate significance below a *p-value* of 0.001 (Bonferroni adjusted) supported by a *chi2* test between predicted and actual classes. The results refer to classifiers which were exclusively trained on *Animal3*, but generalized across animals. Other configurations are shown in the supplemental material (Fig. Sup. 8).

To attribute the effect of stimulation to individual body parts, we trained classifiers on a single variable level. A separate classifier was trained for each animal, burst frequency, and behavioral variable (Fig. 6**b** and Fig. Sup. 8). The rhythmic 10 Hz movements of the *Right Front Paw* is a strong indicator for the 10 Hz stimulation. The height of this paw in the rats’ body reference frame was a highly correlated behavioral variable for the application of the 30 Hz laser stimulation. The pronounced effect on the *Right Front Paw* was in line with the expected outcome for the stimulation of the left motor cortex [20]. More importantly, the classifier exclusively trained on a single sequence of *Animal3* was able to generalize to another sequence of the same animal as well as to recordings of *Animal1* and *Animal2* indicating that FreiPose performs robustly across sessions and animals. Thus, FreiPose allows a detailed comparisons of stimulation effects across animals without the need to retrain for individual cases (Fig. Sup. 8).

## 3 Discussion

We present the marker-free, deep learning based motion capture tool FreiPose for holistic 3D tracking of individual body parts and pose reconstruction of freely moving animals. Instead of triangulating 2D pose reconstructions [8, 9], FreiPose directly reconstructs body poses and movement trajectories in 3D resulting in unprecedented precision. Analyzing the problem holistically by fusing information from all views into a joint 3D reconstruction allows us to surpass the state-of-the-art by 49.4% regarding median error in freely moving rats. We show that FreiPose not only recapitulates previous findings of neuronal correlates of body poses [4], but also allows for automatic clustering of behavior [7] where intuitively meaningful clusters and corresponding variables naturally arise from. As FreiPose has no prior for behavior, no predefinition of behavioral states is needed. This is an advantage over commercial tracking systems which come with a fixed set of detectable behaviors and fail for behaviors outside this scope. Further, we were able to extract single paw trajectories and found astonishingly clear neuronal correlates similar to the ones typically observed in head fixed stereotyped behavioral tasks, e.g. center-out reaching tasks in non-human primates [10, 11]. Importantly, the behavioral clustering and paw trajectories as well as their neuronal correlates can already be extracted from an individual session as short as 30 min. Together, FreiPose directly speaks to the request that behavioral states and uninstructed movements should be taken into account during basically all electrophysiological studies in awake behaving animals [1]. Additionally, analysis of movements can be used as a proxy for monitoring disease state or recovery; e. g. paw trajectories extracted during reaching were used for quantifying stroke recovery [21]. We further demonstrate that optogenetic stimulation effects can be reliably and precisely captured, even reconstructing the temporal dynamics and generalizing across animals and sessions. Given that multiple brain areas have direct projections to spinal cord [22], FreiPose should become a standard tool to objectively quantify or to rule out that optogenetic applications directly evoke an overt motor output or change the frequencies of behavioral categories.

## Supporting information

Video 3 Reaching Rat

Video 4 Behavioral Clusters

Video 1 Rat Pose Tracking

Video 2 Mouse Tracking

## 4 Contributions

C.Z., A.S., T.B. and I.D. conceived and designed the study and wrote the manuscript. C.Z. implemented FreiPose and analyzed the optogenetic data. A.S. conducted electrophysiological recordings, viral injections, fiber implantation as well as optogenetic stimulation and analyzed the neural and behavioral data. M.A. implanted animals with laminar probes.

## 5 Acknowledgements

We thank David Eriksson for setting up the electrophysiology system which was used in this study. We further thank Max Argus, Svenja Melbaum and Philipp Schröppel for proofreading a previous version of the manuscript. This work was supported by the Baden-Wuerttemberg Stiftung, project RatTrack and the cluster of excellence BrainLinks-Brain-Tools (EXC 1086) to T.B. and I.D as well as the Bernstein Award 2012 (01GQ2301) and ERC Starting Grant OptoMotorPath to I.D.

## 6 Online methods

### 6.1 Animals

All animal procedures were approved by the Regierungspräsidium Freiburg, Germany. The animals were housed at a 12 h light dark cycle (light off from 8 a.m. to 8 p.m.) with free access to food (standard lab chow) and water. Three to four animals were pair-housed in type 4 cages (1500U, IVC type 4, Tecniplast, Hohenpeißenberg, Germany) before implantation and single-housed after the implantation in type 3 cages (1291H, IVC type 3, Tecniplast, Hohenpeißenberg, Germany).

#### 6.1.1 Animal surgery

Animals were anesthetized with isoflurane (CP-Pharma, Burgdorf, Germany) inhalation. Subsequently, they were positioned in a stereotactic frame (World Precision Instruments, Sarasota, FL, USA) and their body temperature was kept at 37 °C using a rectal thermometer and a heated pad (FHC, Bowdoin, USA). Anesthesia was maintained with approximately 2 % isoflurane and 1 l/min O_2_. For pre-surgery analgesia, we subcutaneously (s.c.) administered 0.05 mg/kg Buprenorphine (Selectavet Dr. Otto Fischer GmbH, Weyarn/Holzolling, Germany) and Carprofene (Rimadyl, Zoetis, Berlin, Germany). Every other hour, the animals received s.c. injections of 2 ml isotonic saline. Moisturizing ointment was applied to the eyes to prevent them from drying out (Bepanthen, Bayer HealthCare, Leverkusen, Germany). The skin was disinfected with Braunol (B. Braun Melsungen AG, Melsungen, Germany) and Kodan (Schülke, Norderstedt, Germany). To perform the craniotomy, the skin on the head was opened along a 3 cm long incision using a scalpel. The exposed bone was cleaned using a 3 % peroxide solution. Self-tapping skull screws (J.I. Morris Company, Southbridge, MA, USA) for implant stability and for extracellular referencing were placed on the skull. Craniotomies (1×1 mm) were drilled over the prospective implantation sites. For electrophysiological recordings, 3 animals (25-36 weeks) were implanted with 32 channel laminar probes. Those animals were also involved in a different study. In the animal from which the here reported data originated from, we implanted 2 2-shaft silicone probes (E32+R-150-S2-L6-200 NT, Atlas Neuroengineering, Leuven, Belgium) unilaterally in motor cortex (AP: 3.5, ML: 2.5, DV: 2.3 // AP: 0.87, ML: 2.5, DV: 2.26) and connected the probes via a custom interface board to allow the electrophysiological recordings via ZD32 digital headstages (Tucker-Davis-Technologies, Alachua, FL, USA). The craniotomy was sealed with Kwik-Cast (World Precision Instruments, Sarasota, FL, USA) and the implant was fixed using dental cement (Paladur, Kulzer GmbH, Hanau, Germany). The animals were allowed to recover for at least a week.

For the optogenetic perturbation experiments, 3 female Sprague Dawley rats (14 weeks, Charles River, Germany) were unilaterally injected with a ChR2-carrying viral vector (rAAV5/ hSyn-hChR2(H134R)-eYFP-WPREpA, 2.9 × 10^12^ vg/ml, UNC vector core, Chapel Hill, NC, USA) in 2 AP sites with 500 *µ*l each using a 10 *µ*L gas-tight Hamilton syringe (World Precision Instruments, Sarasota, FL, USA). Coordinates corresponded to the caudal forelimb area [23] (Animal 1 - 0.2/0.45 AP, 1.9/2.1 ML, 1.5 DV, Animal 2 - 0.2/0.65 AP, 2/2.1 ML, 1.5 DV, Animal 3 0.3/0.5 AP, 1.85/2.1 ML, 1.5 DV, in mm). Injections were performed at a speed of 100 nl/min, with a 10 min waiting period after each injection. We implanted 200 *µ*m-core fibers (NA 0.39) with metal ferrules (230 *µ*m bore size, both from Thorlabs, Newton, NJ, USA) at a depth of 1.4 mm while avoiding blood vessels. The craniotomies were covered with Kwik-Cast silicone and ferrules secured with super bond (Sun Medical Co., Moriyama City, Japan) and Paladur dental cement. Animals were allowed to recover and express ChR2 for 4 weeks before starting the stimulation.

#### 6.1.2 Optogenetic perturbations

For the connection to the light source (Luxx 473 nm laser, Omicron Laserage, Rodgau, Germany) we employed ceramic sleeves (ADAF1, Thorlabs, Newton, NJ, USA) attached to an elastic spring-suspended custom made patch cord with rotatory joint (Doric Lenses Inc., Quebec, QC, Canada) to allow the rats to freely roam. Rats were stimulated for 5 sec with 30 or 10 Hz square pulses with 10 % duty cycle and 7 mW at the fiber tip with at least 45 sec between stimulation sequences.

#### 6.1.3 Neural recordings

Broadband signals were simultaneously recorded via 2 ZD32 digital head stages connected via a flex-style dual head stage adapter (Intan Technologies LLC, Los Angeles, CA, USA) and electrical commutator (ACO64, Tucker Davis Technologies, USA) to the recording controller (Intan Technologies LLC, Los Angeles, CA, USA). Spike sorting was performed using Mountainsort [24]. Spikes were synchronized to the videos via the camera frame trigger signals. All neuronal data presented in the manuscript originated from one single session of one rat.

### 6.2 Video recordings

The behavioral box for freely moving exploratory behavior was made from 8 mm thick acrylic glass and with a size of 45×36×55 cm. We used either transparent acryl or a metal mesh with 0.7 cm spacing as floor of the box, to also be able to record videos from below.

We used 7 color cameras (6 x acA1300-200uc and 1 x acA800-510uc, Basler AG, Ahrensburg, Germany) placed around the behavioral box to ensure visibility of the rats’ body parts from almost all angles. Ideally, at any time point, the keypoints of interest should be visible from multiple cameras, to ensure accuracy (for more details see Github tutorial). We employed objectives with a focal length 6-8 mm (Kowa, Nagoya, Japan). For calibration of the cameras we employed calibration pattern (described in Sup. 1.2). To ensure planar surface and exact marker size the pattern was manufactured by Schmidt Digitaldruck GmbH, Wörth, Germany.

The cameras recorded with 30 fps using custom python software (see Sup. 1.2). The software is able to set exposure time, gain, frame rate and can adjust the white balance. It allows for individual camera control and recording of calibration sequences. However, FreiPose would work with videos recorded with any other recording systems as long as synchronous frame acquisition is ensured. We used a low cost Arduino based hardware trigger (details and Arduino code available on github) for frame synchronization. The frame trigger signals were also recorded with the Intan recording system to allow for synchronization of the video with neural signals.

- **Tracking of paw during reaching**. We trained a rat in a reaching task. The animal was food-deprived (limited to 3 standard pellets) prior to reaching training. The rat was placed in a transparent box (27×36×29 cm) with an opening (1×6 cm). Sugar pellets were placed in front of the opening. For the reaching task, we used a lens system with larger focal length (12-16 mm, Computar,Cary, NC, USA /Cosmicar, Tokio, Japan) to focus the camera’s image resolution on the reaching movement.
- **Tracking of mice**. We trained FreiPose to predict keypoints on mice. Two C57BL/6 mice from concurrent experiments in our lab were placed in a transparent box (25×35×25 cm), which they explored. The same cameras and objectives were used as in the rat behavioral setup.

### 6.3 Data analysis

#### 6.3.1 1D tuning curves and postural variables

Tuning curves were based on free exploration behavior. In lack of a clearly defined point on the rat’s fur corresponding to its skeletal neck, we used the midpoint between ears as surrogate neck. We defined the body axis of the rat as the vector from the tail root to the midpoint between the ears, and head axis as the vector from the midpoint between ears to the nose. Body and head pitch were calculated as angle between the horizontal and the body or head axis, respectively. Angles between resulting vectors were calculated using trigonometric functions.

To calculate the tuning curves, the head pitch and azimuth were binned in 4 degree steps, body pitch in 2 degree steps. Neural firing rates were binned according to the bin edges (in time) of the body pose variables. By shifting the neural data relative to behavior by ±[5:60] s 1000 times, we obtained a shuffled population. Neurons were considered significantly tuned when their extrema exceeded a z-Score corresponding to a Bonferroni adjusted p-value of 0.05 (quantile function of normal distribution at (1 - 0.05)*/N*) of the shuffled population for at least 3 consecutive postural bins.

For some postural variables (e.g. paw movements) it was required to transform the location of the keypoints to the rat’s local reference frame (such that the rat’s body axis was oriented along the calculated x axis). This distinguishes movements of the paw relative to the body from absolute movements of the paws while the whole body is moving (e.g. during rearing). A rotation matrix was created based on the body axis and the vertical axis (pointing up). Subsequently, keypoints were multiplied with the rotation matrix to obtain keypoints in the new reference frame.

To express movements or postural directions, the direction vector is projected onto a plane (e. g. for the xy-direction of a paw movement, via removing the z-component for the horizontal plane projection). The angle to the ordinate was calculated via basic trigonometry, the histogram of the resulting angles was smoothed with a Gaussian filter for representation purposes and plotted onto a polar projection.

#### 6.3.2 Unsupervised behavioral embedding

We modified an algorithm for behavioral mapping of freely moving fruit flies [13] to cluster the pose reconstruction time-series into behavioral clusters in an unsupervised manner. While the original algorithm relied on postural decomposition via principal component extraction from videos, we employed the FreiPose predicted body keypoints as postural information. Predicted poses were smoothed with a Gaussian (std of 33 ms, 1 time bin) and transformed into the rat body-centric reference frame. Pairwise distances of all keypoints, as well as the distance of each keypoint to the floor plane were combined and the top 20 principal components (explaining 95 % of variance) were extracted. Time courses of the components were Morlet-wavelet transformed with the Py-Wavelets library[25] at the following frequencies: 0.5 and 1-15 Hz in 1 Hz steps. We recombined the temporal features (results of the wavelet transform for all frequencies and components) with the spatial features (keypoint location in the rat’s local reference frame). Combined features were z-Scored and embedded into 2 dimensions via the tSNE approach [26]. We employed the openTSNE python library [27] with cosine distances and PCA initialization, perplexity 100, and exaggeration 12 as parameters. Subsequently, the tSNE embedding was smoothed with a kernel density estimation[28] with a 501×501 dots canvas size and a bandwidth of 0.08. Areas around peak densities were separated via the watershed (scikit-image library) algorithm [29].

For evaluating whether the behavioral clustering resulted in meaningful separations of behavior, time points dwelling in a single cluster for longer than 0.1 s were extracted from the corresponding video and concatenated into a single video file for evaluation by a human observer using ffmpeg (2.8.15) libraries. As t-SNE is a non-parametric approach, the results between run might vary.

Locomotion clusters were typically adjacent in the embedded space between runs, formed a circle or hose-like shape, and were easily identifiable in the videos. Rearing clusters (typically divided into start rearing/rearing/stop rearing clusters) were well separated in the embedded space. Cleaning clusters were characterized by a typical sequence of paw-licking, nose and ears cleaning, and fur cleaning. Further clusters predominately contained periods of movement quiescence with occasional individual steps. Typically this was accompanied with sniffing and whisking and was thus termed ‘exploration of nearby environment’. A more detailed separation of behaviors is possible but is out of scope of this work.

To quantitatively confirm the clustering, we defined intuitive variables, which were expected to discriminate individual clusters. We calculated the paw distance to the nose (expected to be smallest during cleaning), average body pitch (expected to be highest during rearing), horizontal speed of the rat’s center of mass (expected to be highest during locomotion), and the average paw speed (expected to be low during the inspection of nearby environment due to seldom steps, and overall movement quiescence). The difference in the resulting distributions was assessed with the two-sample Kolmogorov–Smirnov test.

#### 6.3.3 Population analysis

Paw velocity in the rat’s local reference frame was thresholded at 0.012 m/frame. This threshold was chosen based on a visual evaluation of speed traces combined with video analysis. The resulted threshold crossings were used for peri-stimulus analysis of the population activity. Neuronal firing rates of each neuron were normalized to have a mean of 0 and a standard deviation of 1. A time window of ±0.5 s around each paw movement onset was extracted from the neural responses as well as from the paw velocity vector. The resulting matrix (neurons × trials × time) was averaged over neurons and trials. Paw velocities were averaged across trials. For comparison with DLC, paw velocity calculated from DLC pose reconstruction was used. For behavioral stratification, neural responses and corresponding paw velocity profiles were indexed according to the corresponding behavioral category during the movement onset. For significance testing, we calculated the mean of the firing rates in a window ±333 ms around the movement onset and compared them across conditions. For the left versus right paw comparison of the neuronal population response we applied a t-test, for individual neurons we used a Bonferroni corrected t-test. For the behavioral stratification of the neuronal population responses, we computed a one-way ANOVA with a post-hoc Bonferroni corrected t-test.

#### 6.3.4 Trajectory stratification

To stratify the front paw movements during freely moving behavior, we extracted periods of high velocity in relation to the body, preceded by periods of lower velocity of the paw defined by a velocity threshold. The threshold (0.012 m/frame) was chosen manually using co-inspection of the velocity trace and the animal behavior. By transferring paw positions into a rat-body-centered reference frame, we extracted paw movements relative to the body. 3D trajectories of the paws were smoothed with a Gaussian (std of 100 ms) and a window ranging from –66 to +500 ms around the timepoint of the threshold crossing was extracted. Trajectories were aligned to start at the origin (0,0,0) of the reference frame. We subdivided the directions of the paw trajectories via k-means clustering (k = 10, using elbow approximation). We stratified the neural responses according to the calculated cluster-labels from the corresponding trajectory. For the raster plots the neuronal spike times were aligned to the frame triggers (and thus the behavior) and segmented into 1.1 ms bins. To obtain continuous firing rates, the spike trains were smoothed with a Gaussian (std of 33 ms). To find neurons which were significantly modulated to different trajectories, we calculated a one-way ANOVA per neuron (dependent variable - sum of spikes in ± [333 ms] window around movement onset, independent variables-trajectory cluster labels) and Bonferroni adjusted the resulting p-values to account for the number of tested neurons. Neurons were considered significantly modulated if the adjusted p-value for the main effect was < 0.05. To define neurons which were significantly modulated by different behavioral states during the execution of similar paw trajectories, we analyzed trajectories corresponding to two similar trajectory clusters (neighboring clusters representing anterior paw movements) to obtain a sufficient number of trajectories. These were stratified depending on the behavioral state during the start of motion. We calculated a one-way ANOVA per neuron on the sum of the spikes in a ± [333 ms] window around movement onset for behavioral categories. p-values were Bonferroni adjusted to account for the number of tested neurons. Neurons were considered significantly modulated if the adjusted p-value was < 0.05.

### 6.4 Motion capture

For motion capture, we combine a bounding box detection network, to extract the region of interest from the full scale images 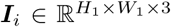, with a novel pose reconstruction architecture (Fig. 7). The approach resolves ambiguities after integration of information from all views. Due to occlusion, it is typically impossible from a single view to measure the exact location of all body keypoints in that view, yet existing methods attempt to reconstruct the key-point locations in the images, regardless of their visibility. This favors learning priors, to hallucinate the invisible keypoints, over measuring their location diminishing performance in the subsequent 3D lifting step. To circumvent the problem, FreiPose extracts features 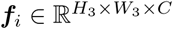 rather than keypoints from the cropped images 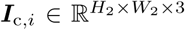 and deploys a differentiable inverse projection operation ∏^*-*1^ [30], which maps features into a 3D representation

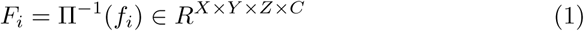

based on the features *f*_*i*_ of camera view *i*. In this notation *H* and *W* represent spatial dimensions and *C* the number of channels for feature representation, which throughout this work are chosen as: *H*_2_ = *W*_2_ = 224, *H*_3_ = *W*_3_ = 28 and *C* = 128. *N* denotes the number of keypoints, which is 12 for freely roaming rodents and 14 in the reaching experiment. Input image resolution *H*_1_ and *W*_1_ lies between 600 and 1280 pixels due to varying image resolutions captured by the cameras deployed. The representations across views are merged by averaging across views ***F*** = 1*/N Σ*_*i*_(***F***_*i*_) and deploy a U-Net-like encoder-decoder architecture 3D CNN [31] on the voxelized representation. The 3D network learns to reason on the joint representation and reconstruct an initial 3D pose ***P*** ^0^ incorporating information from all views. Similarly, to the commonly used map representation for 2D keypoints [32], we use a voxelized representation for localization of each keypoint in 3D. To obtain a point estimate, the location of the voxel with the highest score is retrieved and we call the maximal score the predictions’ confidence *c*_*i*_.

**Figure 7:**
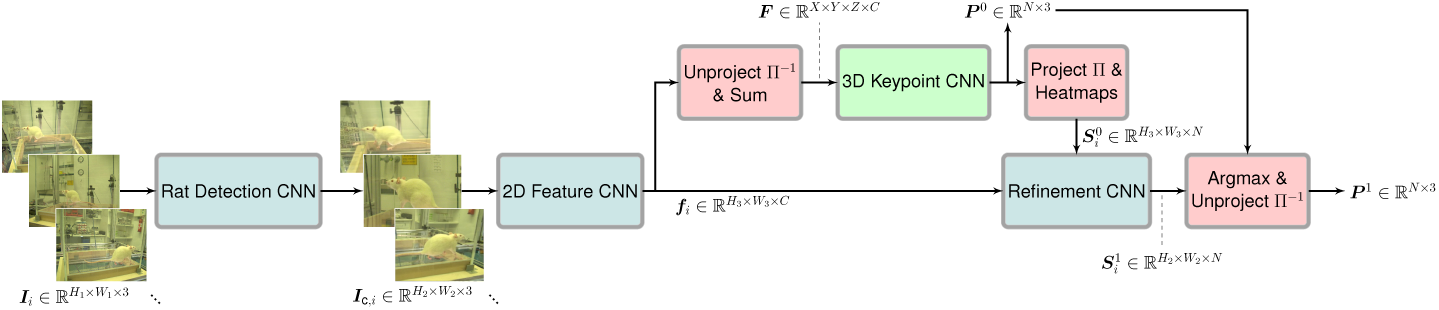
Detailed architecture of the motion capture approach. Given the camera images first a bounding box detection network is applied, then image features are extracted from the cropped images and unprojected into a common 3D representation. The 3D representation is used to reconstruct the initial pose ***P*** ^0^, which is projected into the views for further refinement. Finally, the refined 2D reconstructions 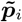 are used to calculate the final 3D pose ***P***^1^. ***S*** is a scoremap representation for a 2D keypoint, which is linked to its point representation through 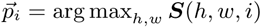 arg max_*h,w*_ ***S***(*h, w, i*) and vice versa through creation of Gaussian target maps [32].

The pose ***P***^0^ ∈ ℝ^*N ×*3^ is a matrix representing the location of the *N* predefined body keypoints at a given time in world coordinates. Subsequently, we use 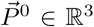 as being a single keypoint sliced from ***P***^0^, or ℝ^4^ in its homogeneous coordinate form if needed. Similarly, 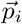 denotes a single 2D keypoint in ℝ^2^ taken from ***p***_*i*_ ∈ ℝ^*N ×*2^ of camera view *i*.

For refinement, the initial 3D pose 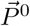 is projected into the camera views

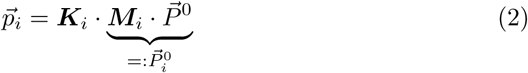

using the cameras’ intrinsic ***K***_*i*_ ∈ ℝ^3*×*3^ and extrinsic matrices ***M***_*i*_ ∈ ℝ^3*×*4^, which are obtained via the camera calibration procedure described in Sup. 1.2.

Given the initial 2D pose 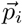 and image features ***f***_*i*_ from view *i*, subsequent convolutional layers estimate refined 2D coordinates 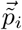. To obtain the final 3D reconstruction 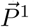 the refined 2D keypoints are unprojected into the world using:

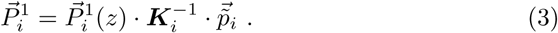

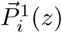 retrieves the third component from the pose in camera coordinates 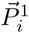, which corresponds to the respective keypoints’ depth in this cameras coordinate frame. Secondly, the scalar reconstruction confidence *c*_*i*_ is used to calculate the final estimate as a confidence weighted average:

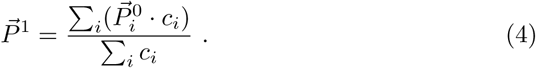

Extensive details on architectural choices and algorithmic hyperparameters are located in the supplemental material or can simply be taken from the released code.

### 6.5 Details of the Motion Capture Experiment

The experimental motion capture results of 2 were obtained by splitting the base dataset of 1199 training and 614 evaluation frames providing 7 cameras into different subsets. For a given number of cameras the experiment (Fig. 1**d**) is repeated in 3 trials choosing different sets of cameras as listed in Table 1. For example, if only 2 cameras are used we pick the following camera pairs: {{1, 5}, {1, 7},{1, 3}}. Each number uniquely identifies a camera (Fig. 1**a**) and the pairs chosen correspond to the cases ‘long side + short side’, ‘long side + bottom’ and ‘long side + long side’. Each of the resulting datasets still covers 1199 time instances, but only 1199 · 2 = 2398 individual frames compared to 1199 · 7 = 8393 in the all camera setting. The same procedure is applied to the evaluation set. Table 1 lists the selected subsets of cameras used for experiments in Fig. 1**d**. Please note, that testing all possible permutations is computationally very expensive, why we resort to testing manually chosen subsets representing meaningful cases, i. e. chose cameras how one would if only a limited number of cameras is available.

**Table 1:** Camera subsets for reduced number of views experiment. Given a number of cameras three different subsets of cameras are selected. Reducing the amount of cameras both leaves less frames for training and increases the difficulty to precisely localize keypoints (Fig. 1**d**). Please note that extensive evaluation of all possible configurations is computationally very expensive, so we resort to using manually selected subsets that reflect reasonable camera placements.

| Number of cameras | 1 | 2 | 3 | 4 | 5 | 6 |
| --- | --- | --- | --- | --- | --- | --- |
| Camera sets | {1} | {1, 5} | {1, 5, 7} | {1, 3, 4, 5} | {1, 3, 4, 5, 7} | {1, 2, 3, 4, 5, 7} |
|  | {5} | {1, 7} | {1, 3, 5} | {1, 3, 4, 7} | {2, 6, 4, 5, 7} | {1, 2, 3, 4, 5, 6} |
|  | {7} | {1, 3} | {4, 5, 7} | {1, 4, 5, 7} | {1, 3, 4, 5, 6} | {1, 2, 4, 5, 6, 7} |

To simulate sparsity of labeled samples (Fig. 1**e**) we use all cameras, but randomly select a subset of 10%, 20%, 30%, 60%, 80% or all time instances. For example, in the 20% case there are 1199 · 0.2 = 239 time instances of 7 cameras in the training set, which results in an effective number of 239 · 7 = 1673 camera frames used. This dataset is used for training both methods, FreiPose and DLC, and evaluation is performed with respect to the complete evaluation set. Each level of sparsity is sampled 3 times for a more robust reconstruction.

Building on the notation introduced in 6.4 the median error is calculated as follows: Let 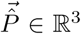 denote the predicted keypoint coordinate of one keypoint and 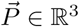 represent the label then the reported metric is defined as

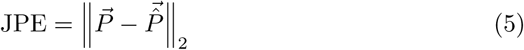

and represents the **J**oint **P**osition **E**rror (JPE). For Fig. 1**d, e** the median, 30% and 70% percentiles of the JPE are calculated over all trials, evaluation frames and keypoints.

### 6.6 Skeletal models

During the freely moving rat experiment we use a 12 keypoint model (Fig. 8**a**), which includes keypoints along the body axes, faces and paws (Fig. 8**b**).

**Figure 8:**
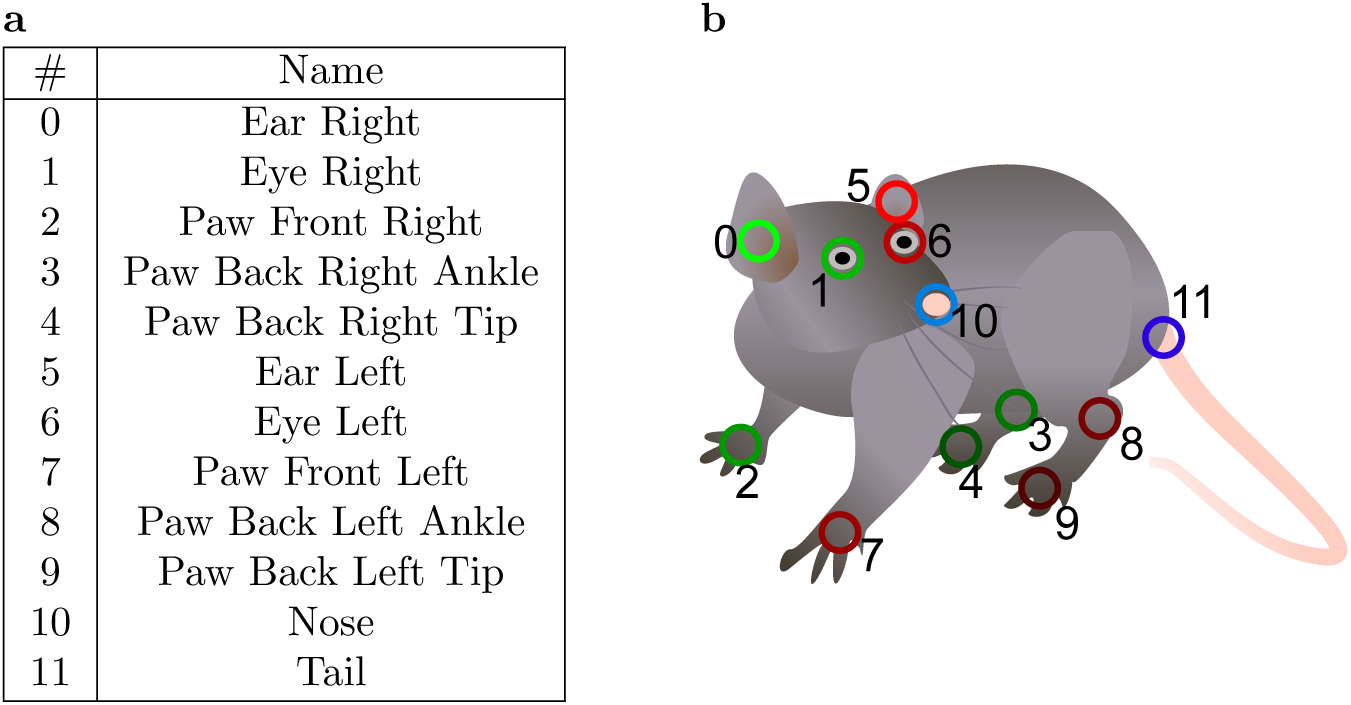
Keypoints defined on the rat body. **a** Names and indices of keypoints. **b** Keypoint locations on the rat body.

For the reaching experiment we defined a 14 keypoint model using a point at the wrist, one for the thumb and three for all the remaining fingers (at metacarpophalangeal, proximal interphalangeal joints and finger tip).

## 7 Data Availability

The raw video data of this study is available from the corresponding author upon reasonable request. Example data sets for FreiCalib and FreiPose are available via our GIN repository: https://gin.g-node.org/optophysiology/FreiPose.

### 7.1 Source data

Source data for reproducing figures of the article will be made available via our GIN repository: https://gin.g-node.org/optophysiology/FreiPose

### 7.2 Code Availability

- Docker runtimes and source code for FreiPose will be made publicly available on Github: https://github.com/lmb-freiburg/FreiPose-docker.
- Docker runtime and source code for FreiCalib is publicly available on Github: https://github.com/lmb-freiburg/FreiCalib.
- Source code for RecordTool will is publicly available on Github: https://github.com/lmb-freiburg/RecordTool.
- **During the review process**, the code of FreiPose is not yet publicly released. Instead, the complete code is located in the Supplemental material as FreiPose-master.zip and FreiPose-docker-master.zip.

## Sup.1 Supplemental material

### Sup.1.1 Experimental setup

The following hardware was used for the experiments presented:

- Cameras: 6 x acA1300-200uc and 1 x acA800-510uc, Basler AG, Ahrensburg, Germany
- Calibration object: Planar aluminium dibond, double sided Apriltag board using 36h11b2 patterns in a 4 × 10 grid, created by FreiCalib and manufactured by Schmidt Digitaldruck GmbH, Wörth, Germany
- Lenses for freely roaming rodents: fixed focal length 6-8 mm, Kowa, Nagoya, Japan
- Lenses for paw reaching: fixed focal lengths of 12-16 mm, by Computar, Cary, NC, USA and Cosmicar, Tokio, Japan
- Camera hardware was mounted to aluminium profiles (ITEM, Solingen, Germany) using custom 3D printed mounts. STL files are available upon request
- Camera mounts: Walimex FT-002H tripod heads
- 2 x Led video lights (Zetci ZT-017, 15 W)
- Experimentation box for freely roaming rats: 45×36×55 cm using 8 mm thick acrylic glass and metal mesh with 0.7 cm spacing
- Experimentation box for freely roaming mice: 25×35×25 cm using 8 mm thick acrylic glass
- Experimentation box for paw reaching: 27×36×29 cm using 8 mm thick acrylic glass and an 1×6 cm opening
- Experimentation boxes were build at the University Freiburg workshop
- Computer specifications: Intel Core i7-6800K, 64 Gb RAM, Geforce GTX 1080, Intel C610/X99 chipset USB controller + Renesas uPD720202 USB3.0 card, Ubuntu 16.04, Python 3.5, Pylon 5.1.0, USB3 kernel Driver xhci hcd
- Electrophysiology: D32 digital headstages (Tucker-Davis-Technologies, Alachua, FL, USA), electrical commutator (ACO64, Tucker Davis Technologies, USA), RHD2000 Evaluation Board 1.0 (Intan technologies, Los Angeles, CA, USA)
- Laser source: Luxx 473 nm laser (Omicron Laserage, Rodgau, Germany) was controlled via QPIDe control board (Quansar,Markham, ON, Canada) from MATLAB R2014 (MathWorks, Natick, MA, USA)

### Sup.1.2 Framework

We introduce a ready-to-use framework for general purpose 3D keypoint estimation from multiple views, which is outlined in Fig. Sup. 1. It is versatile in terms of the number of cameras and their relative positioning as well as the number and semantic meaning of keypoints, which allows application to a broad range of experimental setups.

The framework is easy to install because a *Docker* container is provided in our Github repository https://github.com/zimmerm/FreiPose-docker along with video tutorials that provide a beginner-level introduction about how to use FreiPose for experimentation. We strongly recommend making use of the video tutorials.

**Figure Sup. 1:**
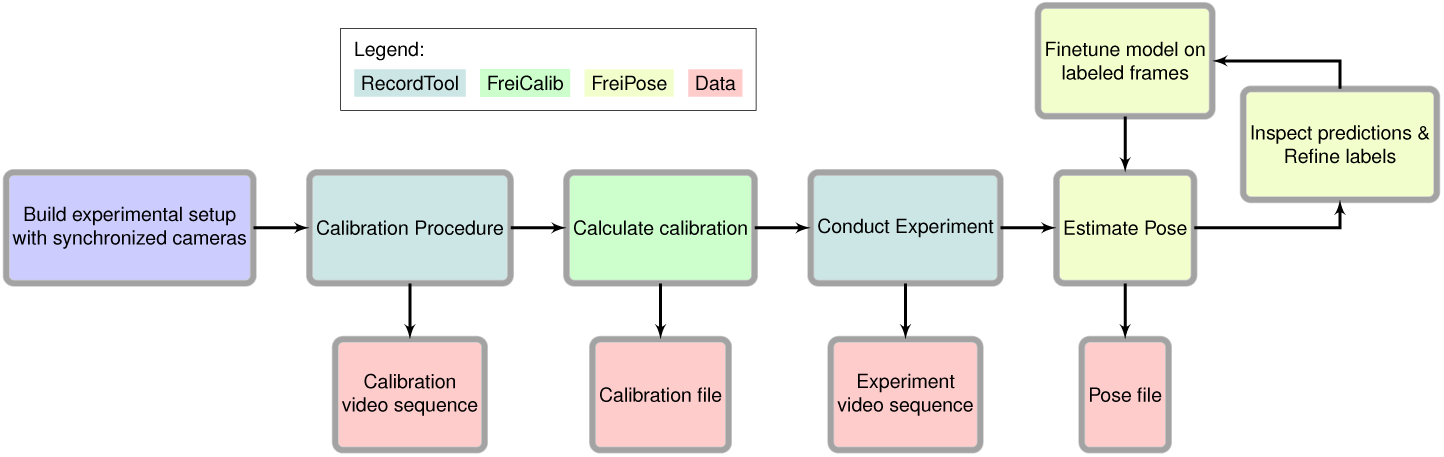
Experimentation work flow. First, the experiment is set up using time synchronized cameras that are calibrated subsequently using FreiCalib and a dedicated calibration recording with RecordTool. Afterwards, experiments can be recorded and the animals’ pose is estimated subsequently. The pose is estimated in a predict-inspect-refine loop that starts with an initial model to make predictions. The predictions are manually inspected and refined if they are not yet sufficiently accurate. The refined predictions are added to the training set to finetune the model. The updated model yields improved predictions. After some iterations, no further manual intervention is necessary and the model yields accurate pose estimates for new recordings.

We divided our framework into three modules, which allow using them independently. The modules are RecordTool, FreiCalib and FreiPose.

- **RecordTool** contains software for recording time synchronized videos. It directly handles the Basler cameras we used in our experiments. When using other cameras, this software must be modified accordingly.
- **FreiCalib** calculates the camera calibration based on time-synchronized video recordings, where an calibration object is used. Notes on how to create such a pattern and hints what makes a good calibration sequence are given in our respective Github repository.
- **FreiPose** estimates poses based on time synchronized videos taken by previously calibrated cameras.

#### Computational requirements

Running FreiPose involves training large neural networks so certain hardware requirements should be met. This involves an Ubuntu 18.04 operating system and a nVIDIA GPU with at least 8GB of memory. Similar operating systems that also provide nVIDIA Docker might work as well, but were not tested. Access to a solid-state drive (SSD) for storing training data is beneficial, but optional.

#### Video Recording with RecordTool

The module **RecordTool** builds upon the proprietary camera drivers of the Basler USB cameras which we used in our study and provides basic functions for camera usage like starting or stopping during recordings as well as modifying relevant camera parameters like white balancing, frame rate, gain, and exposure time. Additionally, the same program operates the Arduino based trigger system.

We designed RecordTool such that it saves the files into one output folder. In this folder, each recording is called a *run* and consists of as many files as there are cameras. The files follow the naming convention `run%03d cam%d.avi`, e.g. `run000 cam1.avi`. The software is available online alongside with a detailed description as pointed out in 7.2.

#### Camera Calibration with FreiCalib

First, video recordings using a suitable calibration object should be obtained. FreiCalib allows creating a PDF file that can be used for manufacturing a pattern calling

~~~
python create_marker.py --tfam 36h11 --nx 10 --ny 4
                        --size 0.05 --double
~~~

which will create two PDF files containing the front and back side of a pattern showing a 10 by 4 grid of tags with size 5 cm each. Alongside with the PDFs (Fig. Sup. 2), a calibration pattern description file is saved that encapsulates marker information for subsequent processing steps.

**Figure Sup. 2:**
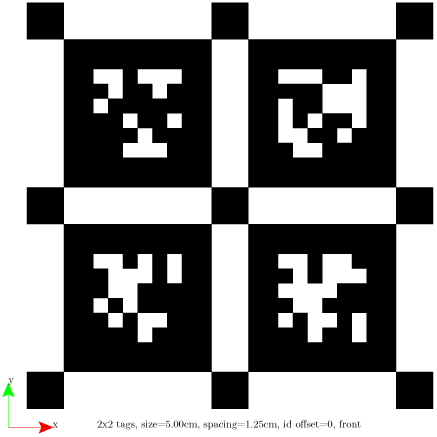
Calibration pattern created by FreiCalib. For simplicity a pattern with *n_x_* = *n_y_* = 2 is depicted. It arranges AprilTags [1] as fiducial identification patterns in a grid structure similar to other calibration methods [2]. Each tag contains a binary code, which uniquely identifies it and the codes are chosen towards robust identification.

For a given recorded calibration sequence, the important parameters describing the imaging process can be calculated by running

~~~
python calib_M.py {MARKER_PATH} {VID_PATH}
~~~

where `{MARKER_PATH}` is the path to the marker description file and `{VID_PATH}` points to the videos showing the calibration sequence. The calibration result will be saved to the folder of the videos as `M.json`. The calibration file contains readable characters and can be opened with any text editor. It contains a dictionary of lists, where each dictionary item presents either extrinsic, intrinsic or distortion parameters and the list iterates over cameras.

Additionally, we provide the option to check a previously calculated calibration with respect to a newly recorded video sequence.

~~~
python check_M.py {MARKER_PATH} {VID_PATH} {CALIB_FILE}
~~~

This allows for easy verification of the given calibrations validity, e. g. none of the cameras was (accidentally) moved or altered between calibration and experimentation time.

#### Experimentation using FreiPose

After obtaining a valid calibration, the actual experiment can be conducted. To get started, an initial configuration of the framework is necessary, which mainly involves defining the keypoints needed for the task and additional task specific settings. A detailed description of these parameters is located in the Github repository and is encapsulated in a configuration file denoted by `{CFG_FILE}`.

To predict poses on a given set of videos, the calls

~~~
python predict_bb.py {CFG_FILE} {VID_PATH}
python predict_pose.py {CFG_FILE} {VID_PATH}
~~~

are sufficient. The commands will save predictions into a new file denoted by {PRED_FILE} in the folder of the videos and its content can be visualized with

~~~
python show_pred.py {CFG_FILE} {PRED_FILE}
~~~

To inspect the predictions in detail and select frames for manual annotation the Selection Tool (Fig. Sup. 3**a**) is used for. It is started by calling

~~~
python select.py {CFG_FILE} {PRED_FILE}
~~~

The user is guided through the selection process by prediction confidences and automatic selection methods presented by the GUI. Frames selected for labeling are extracted from the videos and saved in a separate folder `{FRAME_FOLDER}` as individual frames. The labeling tool (Fig. Sup. 3**b**) can be used in parallel by multiple persons and is started by

~~~
python label.py {CFG_FILE} {FRAME_FOLDER}
~~~

The labeling tool can load the networks’ pose predictions, which allows for speeding up the labeling procedure as the networks’ predictions improve. The user can resort to correcting erroneous labels instead of labeling the complete set of keypoints from scratch. The labeling tool leverages multi-view geometry, i.e. it triangulates the annotations in the individual camera views to a 3D hypothesis and visualizes their locations. This allows the annotator to stop annotating as soon as the hypothesis is consistent across views, which yields a tremendous reduction of labeling effort because usually annotation in few camera views is sufficient.

The folder containing the labeled data is registered to the framework by creating an entry in its configuration file. Afterwards, the network can be fine tuned by calling

~~~
python train_bb.py {CFG_FILE}
python train_pose.py {CFG_FILE}
~~~

**Figure Sup. 3:**
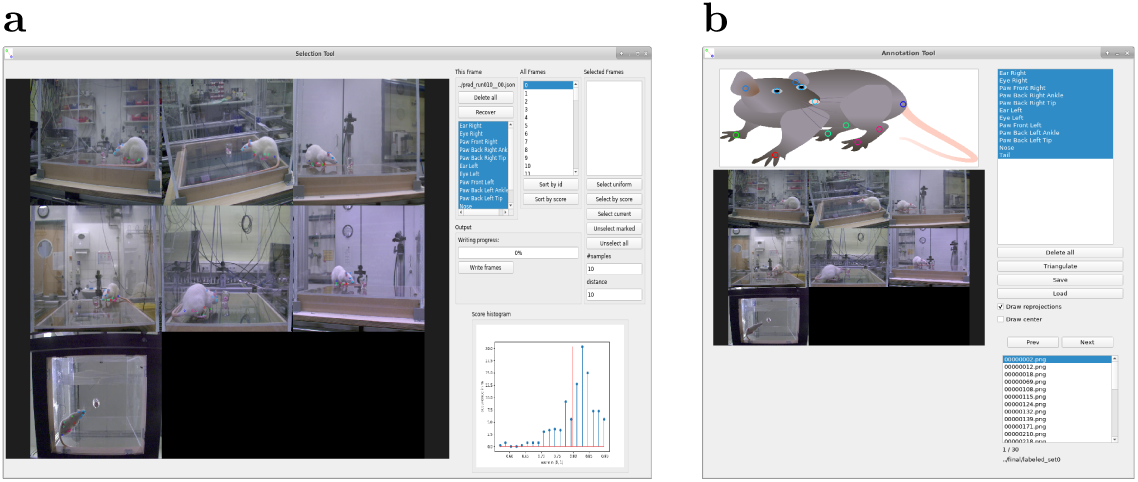
GUIs for viewing predictions and refining annotations. **a** User interface for inspection and selection of frames, where refined annotation is necessary. The selection is guided by a confidence score of the network (lower right corner). **b** The selected frames are extracted from the videos and can be labeled by the annotation tool, which enables the user to annotate by dragging keypoints from the example image and dropping them onto the camera frames. It leverages the multi-view information by calculating a 3D point hypothesis based on the annotated 2D points.

### Sup. 1.3 Additional Experimental Results

We complement the main papers’ experiments with Fig. Sup. 4, which provides JPE results on a per keypoint level in the full dataset and full camera setting. Fig. Sup. 4**a** shows the JPE as box plots for both approaches. Largest errors are present for the highly articulated paw and tail keypoints. Compared to FreiPose the error and variance of DLC is much larger for these keypoints. Fig. Sup. 4**b** is obtained by calculating the percentage of predictions that do not exceed a certain error threshold, which shows that FreiPose can detect keypoints much more reliably than DLC. Within an 5 mm error threshold FreiPose can detect 57.3% keypoints compared to 28.7%, which DLC can detect.

**Figure Sup. 4:**
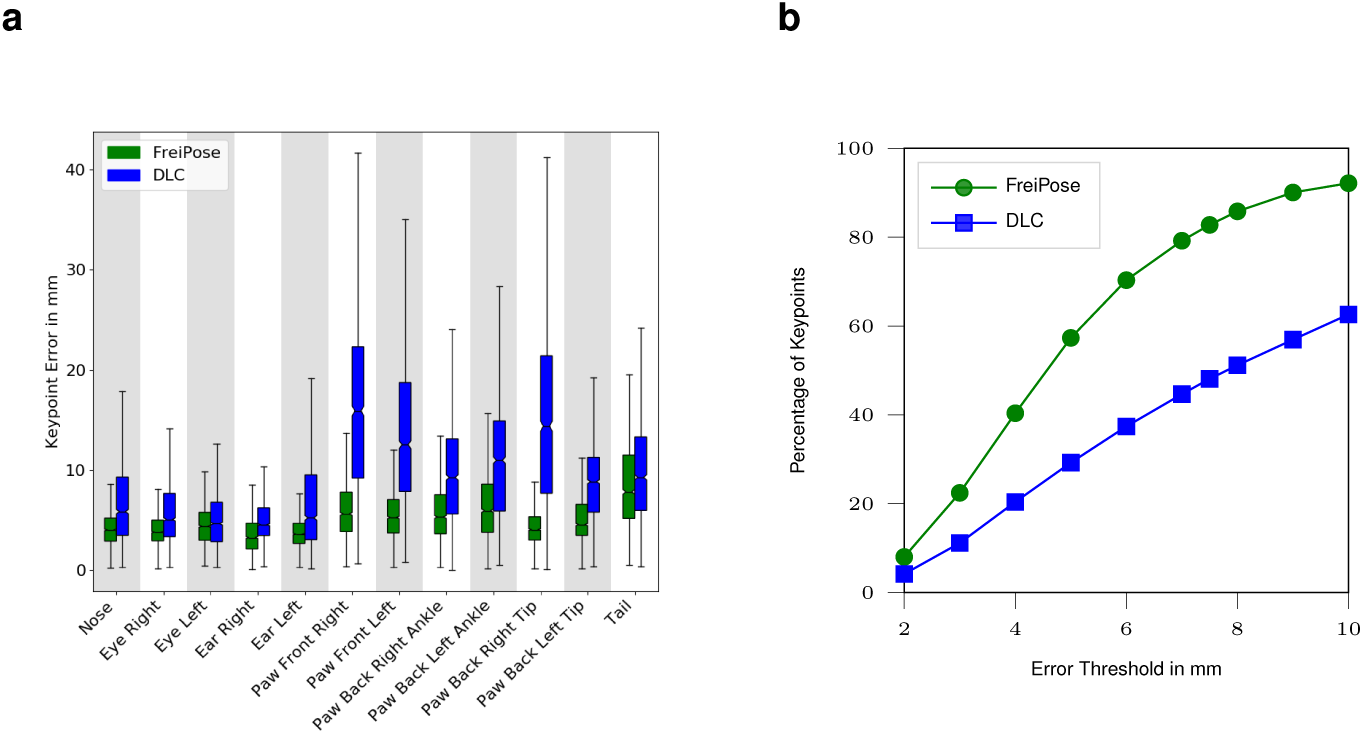
Keypoint errors and percentage of correct keypoints for freely moving rats. **a** Box plot of the keypoint prediction error per keypoint for FreiPose (green) and DLC (blue), where whiskers indicate 1.5 *×* IQR. Largest improvements between FreiPose and DLC are observed for highly articulated keypoints, e. g. paws. **b** Percentage of correct keypoints for a given threshold. For any given error tolerance FreiPose retrieves more keypoints correctly than DLC. Both plots refer to the evaluation set containing 614 samples of three trials.

#### Qualitative Examples

Comparison of DLC and FreiPose method on a qualitative basis shows that the DLC based single view estimation plus post-hoc triangulation is prone to erroneous predictions from individual views (Fig. Sup. 5). DLC was run with a RANSAC based triangulation method to take outlier measurements into account. Keypoint predictions with a confidence below *c* = 0.1 were discarded. The triangulation method is part of the released code within the FreiPose Github repository. Despite these modifications, DLC’s predictions were not reliably correct.

#### Swing-stance classification from pose prediction

We used FreiPose to classify the paw states into swing and stance phases for each of the 4 paws individually, by adding a simple classifier. The classification was based on behavior variables describing Paw Height over Ground and the Paw Velocity, which are calculated from the pose estimation. We use a linear SVM trained on 79 frames and evaluated on 96 different frames of the same animal. Comparison to a manually labeled ground truth shows, that the classifiers trained on our prediction reaches a balanced accuracy of 79.1% compared to 58.2% for DLC averaged over all paws. This shows that increased accuracy of the pose estimation is important for downstream tasks. Detailed results are summarized in Table Sup. 1.

**Table Sup. 1:**
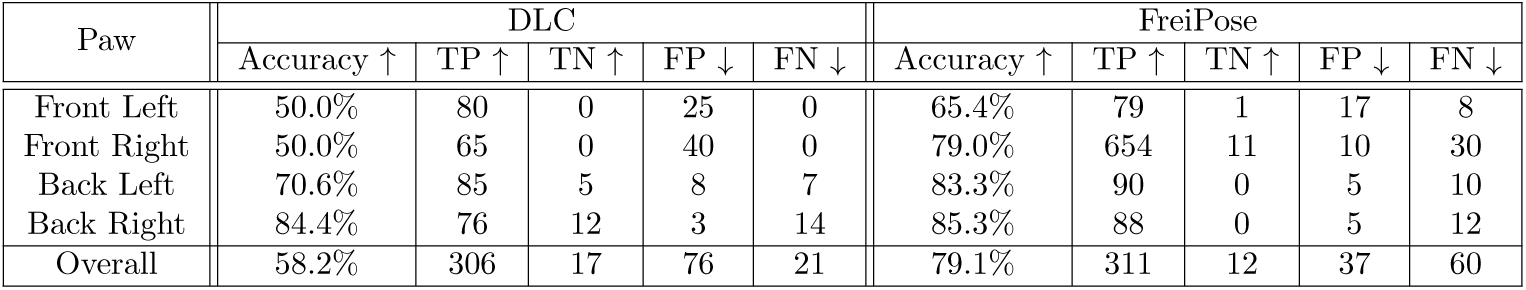
Extended result of swing stance classification. Positive indicates presence of *swing* state of the respective paw. For calculating the accuracy we use the balanced class weight average to account for the imbalance in number of samples of both classes, i. e. (*tp* + *tn*)*/*(*tp* + *tn* + *fp* + *tn*). tp - true positive, fp -false positive, tn-true negative, fn-false negative.

| Paw | DLC |  |  |  |  | FreiPose |  |  |  |  |
| --- | --- | --- | --- | --- | --- | --- | --- | --- | --- | --- |
| | Accuracy ↑ | TP ↑ | TN ↑ | FP ↓ | FN ↓ | Accuracy ↑ | TP ↑ | TN ↑ | FP ↓ | FN ↓ |
| Front Left | 50.0% | 80 | 0 | 25 | 0 | 65.4% | 79 | 1 | 17 | 8 |
| Front Right | 50.0% | 65 | 0 | 40 | 0 | 79.0% | 654 | 11 | 10 | 30 |
| Back Left | 70.6% | 85 | 5 | 8 | 7 | 83.3% | 90 | 0 | 5 | 10 |
| Back Right | 84.4% | 76 | 12 | 3 | 14 | 85.3% | 88 | 0 | 5 | 12 |
| Overall | 58.2% | 306 | 17 | 76 | 21 | 79.1% | 311 | 12 | 37 | 60 |

**Figure Sup. 5:**
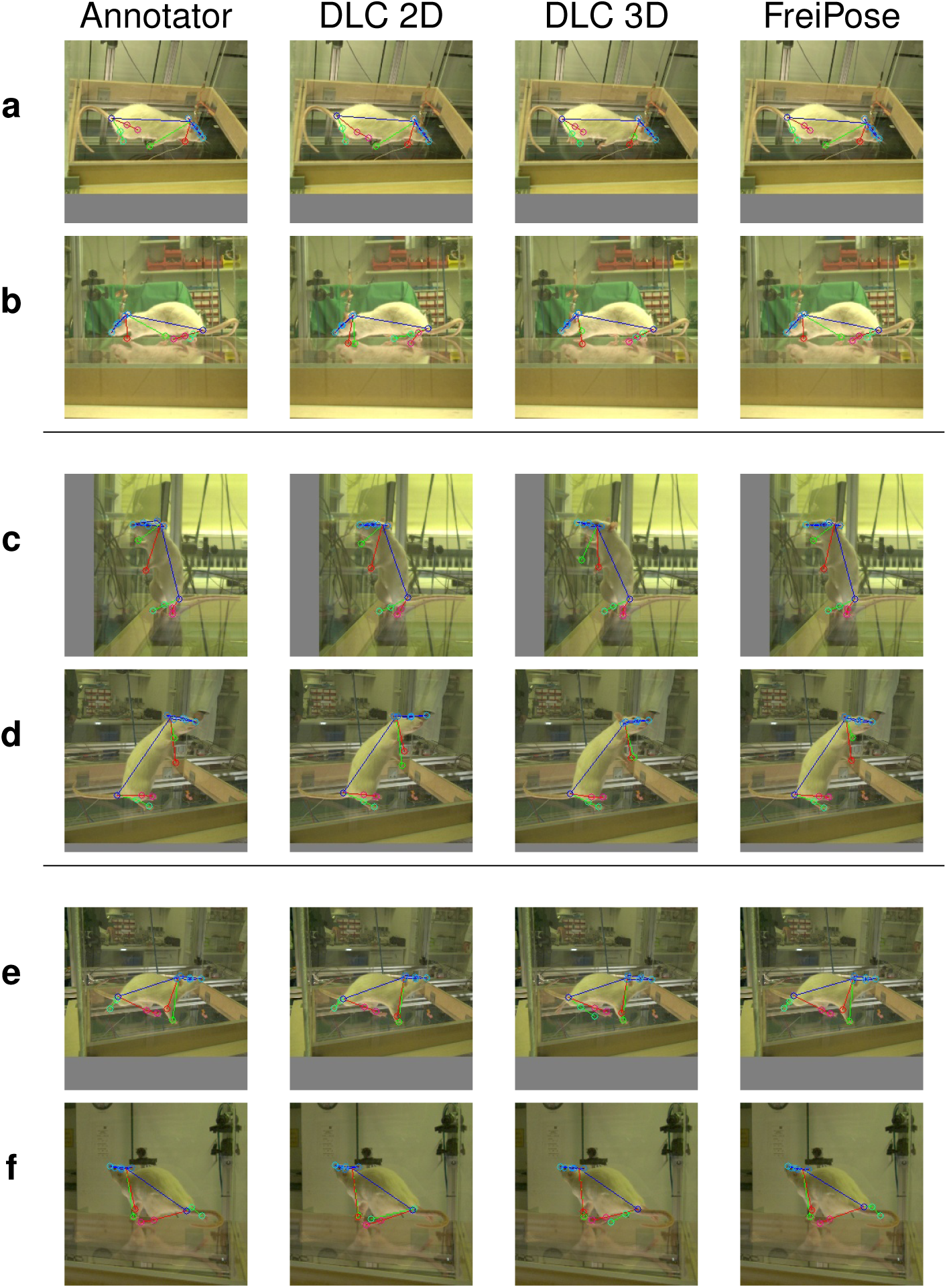
Qualitative comparison between FreiPose and DLC. Rows **a, b**, respectively **c, d** and **e, f**, show images recorded at the same time but from different cameras. DLC is able to correctly estimate poses in rows **a, c, e** but during triangulation from 2D predictions to 3D points the less accurate predictions from **b, d** and **f** have a deteriorating influence on the final results even though robust triangulation techniques were used. On the other side FreiPoses’ predictions look visually similar to the annotations created by a human annotator.

#### Behavioral Embedding

In the main article (Fig. 3) we clustered the movements to identify time points corresponding to 4 types of movement related behavior: locomotion, rearing, cleaning, and inspection of nearby environment. Here we provide an overview of the clustering procedure (Fig. Sup. 6). A more detailed description is part of the online methods (6.3.2).

**Figure Sup. 6:**
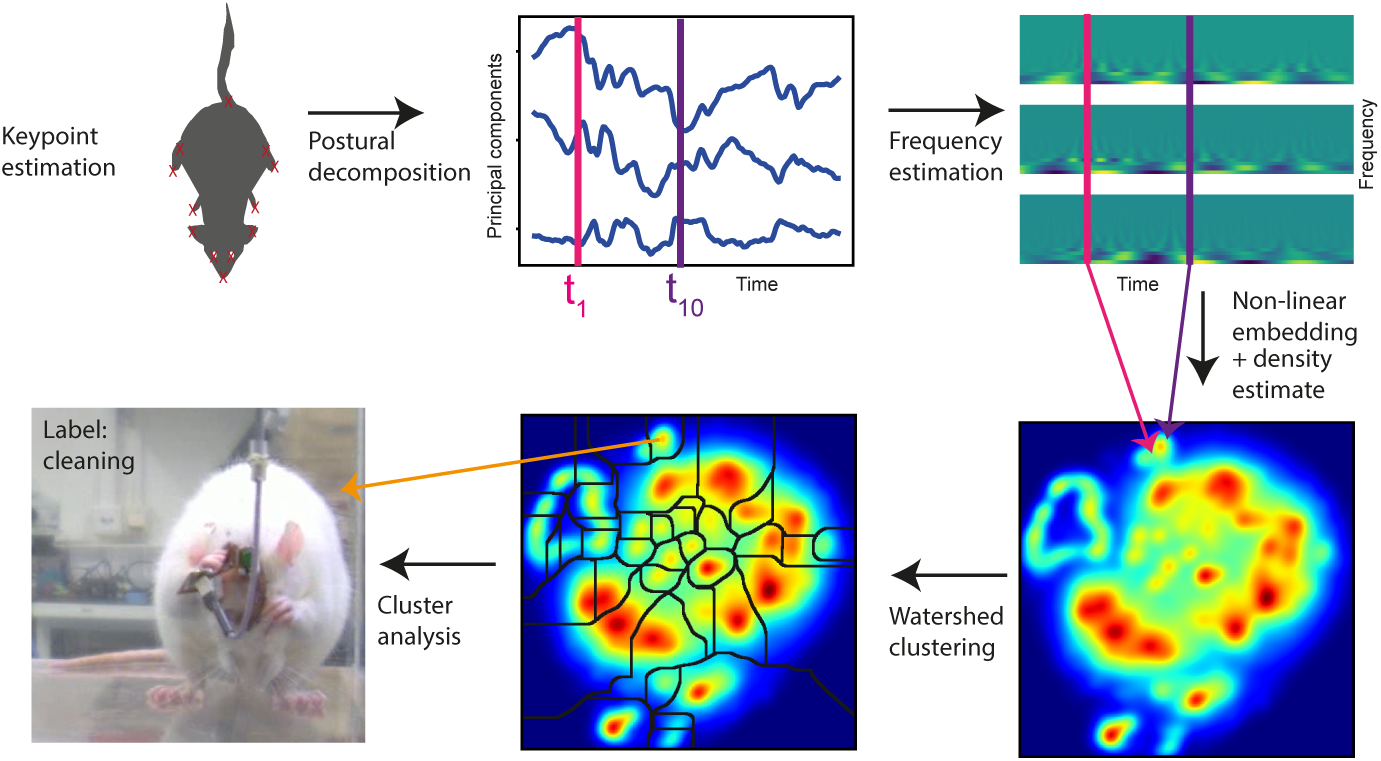
Overview over behavioral clustering algorithm. The modified algorithm for unsupervised clustering of behavior based on a mapping approach for fruit flies [3]. Postural decomposition was achieved via PCA of the pairwise-distance matrix of keypoints. The frequency of the components was calculated via wavelet transform. Extracted features were embedded via t-SNE and individual behavioral clusters identified with watershed clustering followed by assignment of clusters to behavioral labels by visual inspection and evaluation. Refers to main Fig. 3

#### Trajectory Clustering

In the main article in Fig. 4 and Fig. 5 we showed the application of FreiPose to detect modulation of neurons due to paw movements. We applied the same analysis based on DLC (Fig. Sup. 7). The detection of neuronal modulation with DLC is very limited in freely moving animals.

**Figure Sup. 7:**
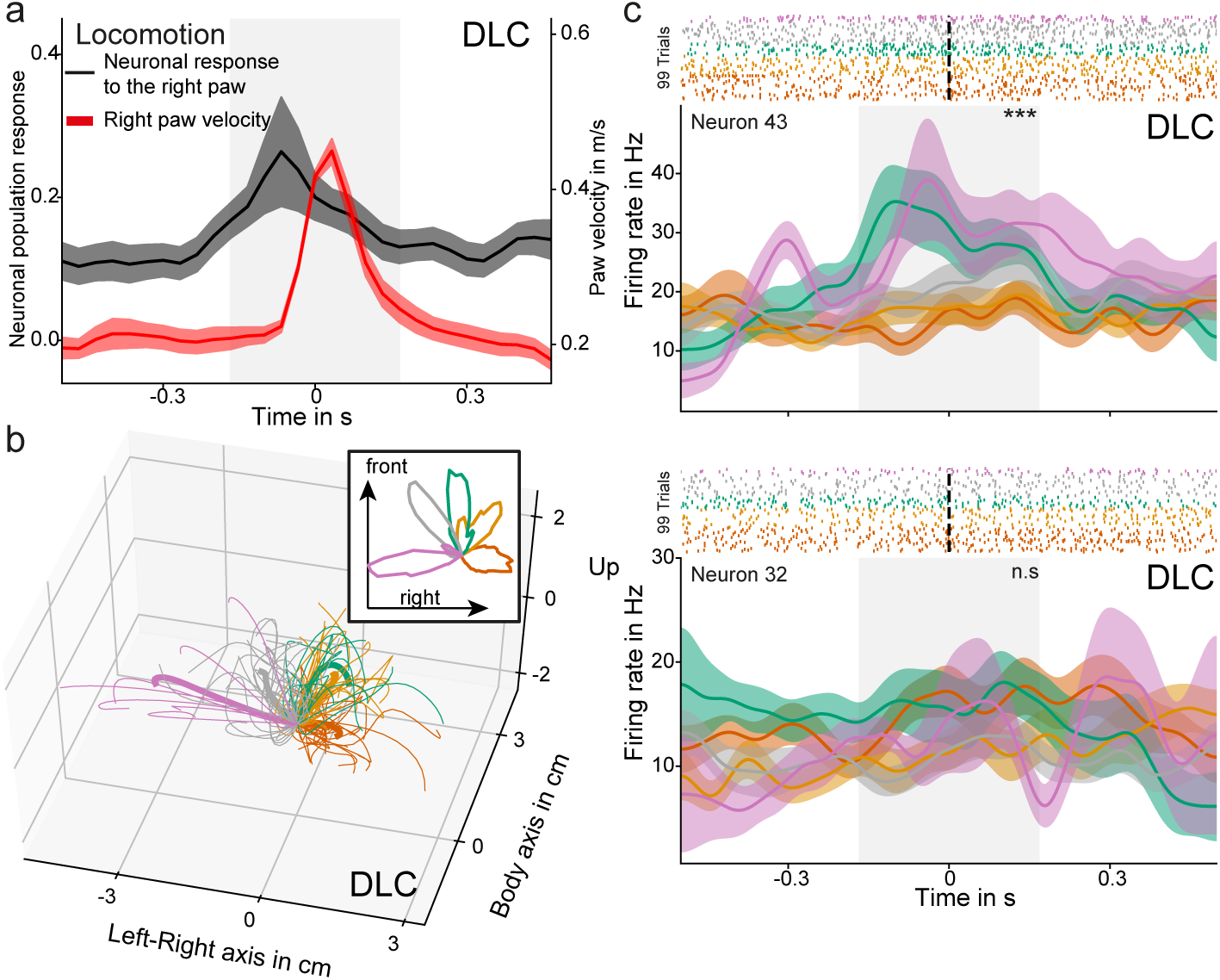
DLC predictions impair the detection of neuronal tuning compared to holistic 3D prediction from FreiPose. **a** Population response to right (contralateral) paw movements during locomotion. **b** Trajectories of the right paw during step-like paw movements extracted via DLC predictions. Note the strong bias to the left, putatively originated from the tendency of DLC to miss-label left and right paws. **c** The responses of the two neurons from main Fig. 5 have been reanalyzed based on DLC extracted paw trajectories. Similar tuning for neuron 43 could be extracted. However, no tuning was detected based on DLC predictions for neuron 32. Refers to main Fig. 4 and Fig. 5

#### Optogenetic Stimulation Experiment

In the main paper, we showed that a classifier trained on Animal3 generalizes consistently to the remaining animals (Fig. 6). We obtained the same generalization when the classifier was trained on Animal1 (Fig. Sup. 8**a**) or Animal2 (Fig. Sup. 8**b**). In addition to the behavioral variables found in Fig. 6 (‘Paw Front Right: Height’ for 30 Hz and ‘Paw Front Right: Height over Ground’), also other variables were detected. Most of the additional variables in the 30 Hz case were related to facial factors that indicate a significant ‘Head Roll’. For classifiers trained on Animal1 or Animal2 similar periodic movement relative to the head is detectable in the 10 Hz case.

**Figure Sup. 8:**
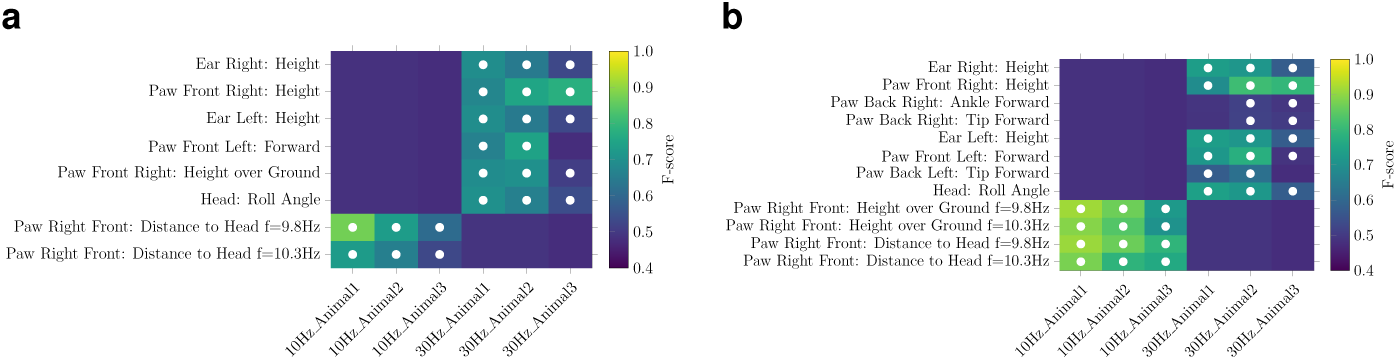
Permutations of the optogenetic stimulation experiment. Similar findings during automatic attribution of the simulation effect when training was performed on Animal1 (**a**) or Animal2 (**b**). Shown is the *F-score* of the respective classifier and white dots indicate significance below a *p-value* of 0.001 (Bonferroni adjusted) supported by the *chi2* test between predicted and actual classes.

### Sup. 1.4 Architectural Details of FreiPose

The implementation of our approach uses Tensorflow [4] and is available through our Github repository.

For bounding box detection we used a COCO [5] pretrained MobileNet V2 [6], which was retrained for the task of detecting the foreground objects. In the freely moving rat scenario, it was trained using each view of the 1199 labeled time instances separately, i.e., a total of 9592 samples. We trained it for 150 k iterations using a learning rate of 0.004 and the RMSProp optimizer. As data augmentation operations, we employed random flipping, cropping, scaling, and color space variation.

For the reaching task, due to the action always happening in the same space, we used a fixed, predefined region of interest rather than a detector.

For pose estimation we extend the approach by Zimmermann et al.[7] by the refinement module for increased accuracy. Detailed discussion of this approach is provided in section 6.4. The network was trained for 60 k using ADAM optimizer [8] with a base learning rate of 10^*-*4^ and decay by a factor of 0.1 every 30 k steps. To improve convergence we found it helpful to not train the refinement module for the first 30 k steps.

**Table Sup. 2:** Tested behavioral variables. ∢ (·) measures the angle between two three dimensional vectors and [*a, b*] defines an vector that goes from point *a* to point The notation {*a, b*} calculates the average of the points *a* and *b*. 3D points are denoted as keypoint indices, that find their textual counterpart in Table 8. The rat local coordinate frame is defined with its origin at {0, 5}, its z axis being aligned with the up pointing normal of the ground plane and its y axis rotated towards [{0, 5}, 11]. STFT - short-time Fourier transform.

| Name | Number of factors |
| --- | --- |
| Keypoints in cartesian rat local frame | $12 \cdot 3 = 36$ |
| Keypoints velocity in cartesian rat local frame | $12 \cdot 3 = 36$ |
| Keypoints distance to rat local frame origin | 12 |
| Keypoints distance velocity to rat local frame origin | 12 |
| STFT of Keypoint distance to rat local frame origin using 33 frequencies | $12 \cdot 33 = 396$ |
| Keypoint distance to ground plane | 12 |
| STFT of Keypoint distance to ground plane using 33 frequencies | $12 \cdot 33 = 396$ |
| Body elevation: $\angle(g, [11, \{0, 5\}])$ | 1 |
| Head elevation: $\angle(g, [10, \{0, 5\}])$ | 1 |
| Head roll: $\angle(g, [0, 5])$ | 1 |
| Head nick: $\angle([11, \{0, 5\}], [\{0, 5\}, 2])$ | 1 |
| Paw right: $\angle([11, \{0, 5\}], [\{0, 5\}, 2])$ | 1 |
| Paw left: $\angle([11, \{0, 5\}], [\{0, 5\}, 7])$ | 1 |
| Angle velocity of the aforementioned angles | 6 |
| Total | 912 |

